# Remodeling oligodendrocyte lipid metabolism via liver X receptors overcomes inflammatory blockade of remyelination

**DOI:** 10.64898/2026.08.07.743529

**Authors:** Judy J. Lee, Matthew D. Smith, Xiaojing Deng, Jingwen Hu, Andrew Love, Jerry S. Jing, Payam Gharibani, Pragney Deme, Abdulshakour Mohammadnia, Qiao-Ling Cui, Daryan Chitsaz, Asmita Dhukhwa, Jaime Gonzalez Cardona, Kathryn C. Fitzgerald, Cole A. Harrington, Xitiz Chamling, Jack P. Antel, Norman J. Haughey, Peter A. Calabresi, Michael D. Kornberg

**Author notes:** Correspondence to: Michael D. Kornberg Johns Hopkins School of Medicine, 1729 East Monument Street, 5 North – 5326 Baltimore, MD, USA 21287.

## Abstract

Multiple sclerosis is characterized by immune-mediated demyelination and inefficient remyelination, owing to impaired differentiation of oligodendrocyte precursor cells (OPCs) into myelinating oligodendrocytes (OLs). Inflammatory cytokines within multiple sclerosis lesions inhibit OPC maturation and induce an immune-like phenotype with antigen-presenting properties, but the underlying mechanisms remain poorly defined. Here, we show that inflammation reprograms OPC lipid metabolism, linking altered metabolism to remyelination failure. In cultured rodent OPCs, interferon-γ (IFN-γ) induced a switch from lipid synthesis to utilization, leading to reduced intracellular fatty acid levels and increased dependence on fatty acid oxidation. Transcriptional analyses confirmed similar lipid metabolic changes in OL-lineage cells cultured from human surgical specimens or isolated from mouse models of inflammatory demyelination and human multiple sclerosis lesions. Enhancing lipid availability in OPCs through oleic acid supplementation or inhibition of fatty acid oxidation attenuated immune-like functions and increased differentiation. Pharmacologic activation of liver X receptor (LXR) transcription factors rebalanced lipid metabolism, suppressed immune-like functions, and overcame IFN-γ–induced differentiation blockade in both mouse and human-derived OPCs. In an adoptive transfer-cuprizone mouse model in which inflammation directly impairs remyelination, LXR activation increased mature OL generation and augmented myelin repair. Together, these findings identify lipid metabolic remodeling as a key mechanism by which inflammation impairs OPC differentiation and highlight LXR activation as a therapeutic approach to enhance remyelination in multiple sclerosis.

## Introduction

Multiple sclerosis is a chronic inflammatory disease of the central nervous system (CNS) characterized by immune-mediated demyelination and loss of oligodendrocytes (OLs).^1,2^ Chronic demyelination leads to progressive neurologic disability by both disrupting axonal conduction and depriving neurons of myelin-derived metabolic and trophic support.^3–6^ Current immunomodulatory therapies target the peripheral immune system to limit clinical relapses and new lesion formation but do not restore myelin, leaving remyelination failure a major unmet clinical challenge.^7^

Oligodendrocyte precursor cells (OPCs) retain the capacity to regenerate myelin through differentiation into mature OLs (mOLs). Although OPCs remain abundant in multiple sclerosis, differentiation into mOLs appears to be blocked in many lesions, leading to remyelination failure.^8,9^ Accumulating evidence indicates that inhibitory factors within the lesion microenvironment play a major role in preventing OPC differentiation, with chronic inflammation representing one of the most potent of these factors.^10–15^ Thus, while many current approaches seek to enhance the intrinsic differentiation potential of OPCs, successful remyelinating therapies must also overcome the suppressive effects of chronic inflammation. To this point, treatments such as benztropine, miconazole, and clemastine, which augment OPC differentiation under non-inflammatory conditions, nonetheless fail to overcome cytokine-mediated differentiation blockade.^15^

Inflammatory cytokines not only block OPC differentiation but also induce a distinct phenotype of OL-lineage cells with immune-like characteristics, known as immune oligodendroglia (iOLGs).^16–19^ These iOLGs, which are found both in human multiple sclerosis lesions and in mouse models,^19–23^ are capable of phagocytosis and cross-presentation of antigen via MHC class I and II molecules, leading to T cell activation.^16,24^ Thus, OPCs can adopt divergent fates following inflammatory demyelination, actively participating in inflammation or regenerating myelin through differentiation into mOLs. Identifying the pathways that control these distinct fates is crucial for developing therapies that promote regeneration in an inflammatory context.

Because cellular metabolism critically regulates differentiation (including OPC differentiation)^25–30^ as well as the fate and effector functions of conventional immune cells,^31,32^ we investigated how inflammatory cues alter OPC metabolism. Using a combination of metabolomics and transcriptomics, we first observed that interferon-gamma (IFN-γ), a ubiquitous cytokine within multiple sclerosis lesions^33–37^ that potently inhibits OPC differentiation^11^ and induces iOLG functions,^16^ produces a concerted remodeling of OPC lipid metabolism. This metabolic reprogramming was broadly characterized as a transition from lipid biosynthesis to utilization, leading to depletion of free fatty acids (FAs). By examining existing single-cell transcriptional databases, we observed similar remodeling of lipid metabolism in disease-associated OL-lineage cells in mouse models and human multiple sclerosis lesions, suggesting broad disease relevance. Multiple targeted approaches to modulate lipid metabolism overcame the effects of inflammation on OPCs, preventing iOLG generation and promoting mOL differentiation. Our studies identified the liver X receptor (LXR) transcription factors as master regulators of OPC lipid metabolism that can be therapeutically targeted to enhance remyelination within inflammatory environments *in vivo*. Together, these findings identify LXRs, and lipid metabolism more broadly, as therapeutic targets for overcoming the inflammatory microenvironment to promote remyelination in multiple sclerosis.

## Materials and methods

### Study design

The aim of this study was to investigate the metabolic changes in OPCs induced by inflammatory stimuli in order to identify therapeutic targets for overcoming inflammatory blockade of OPC maturation and enhancing remyelination in multiple sclerosis. To accomplish this aim, we used cultured rodent- and human-derived OL-lineage cells and the adoptive transfer-cuprizone mouse model, combined with analysis of published transcriptional datasets from human cells, mouse models, and human multiple sclerosis tissue. Assay measurements and end points were all prospectively selected. Experimental design and the sample sets from which data were derived are described in detail in the figure legends and the Materials and Methods section. For *in vivo* studies, age- and sex-matched mice were randomized into treatment groups. Analyses were performed by investigators blinded to treatment groups. Power analyses were not performed. All analyses conducted in the manuscript included all data points, without removal of outliers.

### Animals

Rodents were housed and maintained in a pathogen-free animal facility at Johns Hopkins University. All protocols were approved by the Johns Hopkins Institutional Animal Care and Use Committee. OVA257–264 TCR transgenic OT-I mice (C57BL/6-Tg(TcraTcrb)1100Mjb/J, Strain 003831), 2D2 MOG35–55 TCR transgenic mice (C57BL/6-Tg(Tcra2D2,Tcrb2D2)1Kuch/J, Strain 006912) and wild-type C57BL/6 mice (Strain 000664) were purchased from The Jackson Laboratory. Sprague-Dawley rats were purchased from Charles River Laboratories. LXRα^-/-38^ and LXRβ^-/-39^ mice were a kind gift Ira Schulman (University of Virginia) and originally provided by David Mangelsdorf (University of Texas Southwestern Medical Center, Dallas, TX). Single LXR KO mice were bred together at Johns Hopkins to generate LXR double KO mice.

### Isolation and treatment of mouse OPCs

Cerebral cortices from P5-7 mouse pups were dissected in ice-cold HBSS^-/-^ and triturated into single-cell suspension with dissociation buffer 20 U/mL papain (Worthington Biochemical) and 100 U/mL DNase I (Worthington Biochemical). Tissue dissociation was stopped by addition of HBSS^+/+^ and cells were filtered through a 100 µm strainer prior to centrifugation. The cell pellet was resuspended in 0.2% BSA/HBSS^+/+^ and OPCs were isolated by immunopanning as previously described.^40^ Endothelial cells and microglia were depleted with plates coated with Bandeiraea Simplicifolia Lectin-1 (BSL-1, Vector Laboratories) and CD11b (Bio-Rad) antibodies. OPCs were gathered on plates coated with platelet-derived growth factor receptor alpha (PDGFRα) antibodies (BD Biosciences). Plates were washed with HBSS^+/+^ and OPCs were detached from the plate with Trypsin. Cells were centrifuged and seeded on cover slips or plates coated with poly d-lysine (PDL, Sigma). OPCs were maintained in OPC base media supplemented with 20 ng/mL PDGF-AA (Peprotech, 100-13A), 10 ng/mL CNTF (Peprotech, 450-13), and 1 ng/mL NT3 (Peprotech, 450-03) for 48 h. OPC differentiation was induced by changing to media containing 40 ng/mL T3 (Sigma, T6397). 50 ng/mL IFN-γ (Peprotech, 315-05) was added to induce iOLGs. OPC base media was made with Dulbecco’s Modified Eagle Medium containing 4 mM L-glutamine and 1 mM sodium pyruvate (Gibco, 11995-073), supplemented with 2% serum-free B27 (Gibco, 17504044), 100 U/mL penicillin/streptomycin (Quality Biological, 120-095-721), 0.1% trace elements B (Corning, 25-022-CI), 1x SATO [100 μg/mL apo-transferrin (Sigma, T1147), 100 μg/mL bovine serum albumin (Sigma, A4161), 16 μg/mL putrescine (Sigma, P7505), 60 ng/mL progesterone (Sigma, P8783), 40 ng/mL sodium selenite (Sigma, S5261) in Neurobasal medium (Gibco, 21103049)], 10 ng/mL biotin (Sigma, B4639), 5 μg/mL insulin (Sigma, I6634), 5 μg/mL N-acetyl cysteine (Sigma, A9165), and 4.2 µg/mL forskolin (Sigma, F6886).

All treatments such as T3, IFN-γ, and drugs were added at the same time except for A922500. Cells were pre-treated with A922500 for 2 h before IFN-γ treatment. Reagent information is documented in supplemental file S1.

### Isolation and treatment of rat OPCs

Neonatal Sprague-Dawley rat pups (P2–P5) were euthanized by decapitation, and brains were removed from the skulls. Cortices were manually dissected, minced into fine pieces, and dissociated using the Neural Tissue Dissociation Kit – Papain (Miltenyi, 130-094-802) according to the manufacturer’s recommended protocol. OPCs were enriched by A2B5-positive selection (Miltenyi, 130-093-392) according to the manufacturer’s recommendations. To minimize microglial contamination in subsequent cultures, positively selected cells were stained with anti-CD45-APC-eFluor780 (1:100, eBioscience, 47-0461-80) and anti-A2B5-PE (1:20, Miltenyi, 130-123-715) for 30 minutes in MACS buffer (0.05% BSA, 2mM EDTA in PBS), then washed in MACS buffer and CD45-;A2B5+ cells were sorted on a FACSAria II prior to plating.

Sorted cells were cultured for 4 days under proliferative conditions as previously described.^41^ Briefly, cells were cultured in Dulbecco’s Modified Eagle Medium containing 4 mM L-glutamine and 1 mM sodium pyruvate (Gibco, 11995-073), supplemented with 2% serum-free B27 (Gibco, 17504044), 100 U/mL penicillin/streptomycin (Quality Biological, 120-095-721), 0.1% trace elements B (Corning, 25-022-CI), 100 μg/mL apo-transferrin (Sigma, T1147), 100 μg/mL bovine serum albumin (Sigma, A4161), 16 μg/mL putrescine (Sigma, P7505), 60 ng/mL progesterone (Sigma, P8783), 40 ng/mL sodium selenite (Sigma, S5261), 10 ng/mL biotin (Sigma, B4639), 5 μg/mL insulin (Sigma, I6634), 50 ng/mL hydrocortisone (Sigma, H0135), 5 μg/mL N-acetyl cysteine (Sigma, A9165), and 20 ng/mL recombinant PDGF-AA (PeproTech, 100-13A).

After 4 days in culture, one set of wells from each biological replicate was collected as the baseline time point. The remaining wells were refreshed with differentiation medium consisting of the same formulation as the proliferation medium, but supplemented with 10 nM T3 (Sigma, T6397), in the presence or absence of 10 ng/mL IFN-γ. Cells were stimulated for 24 h.

### iOLG – CD8^+^ T cell co-culture

iOLG-CD8^+^ T cell co-cultures were performed as previously described.^40^ Briefly, mouse OPCs were treated with IFN-γ for 16 h and then spiked with whole Ovalbumin protein (Sigma) for 8 h. Cells were washed twice with PBS to remove cytokines and unprocessed ovalbumin. CD8^+^ T cells were isolated from OT-I mice with CD8^+^ isolation kit (BioLegend) and were added at a 3:1 CD8:OPC ratio for 24 h. Protein Transport Inhibitor Cocktail (eBioscience) was added 8 h before harvesting the CD8^+^ T cells.

### Differentiation and culture of genome-engineered hPSC-derived OPCs

Differentiation of hPSCs was performed as previously described.^42^ Briefly, using CRISPR-Cas9, super-fold GFP (sfGFP) and Secreted Nano-luciferase (secNluc) reporter sequences were knocked-in before the stop codons of the PLP1 and MBP genes, respectively, into a previously engineered hESC reporter cell line (PDTT)^43^ that enables purification of PDGFRA expressing oligodendrocyte lineage cells with identification-and-purification (IAP) tag. This triple reporter cell line allows PDGFRA expressing OPCs to be purified after hESC differentiation, and quantification of mature oligodendrocyte efficiency by secNluc expression on a time course. These cells were differentiated for 75 days to generate OPCs. OPCs expressing PDGFRα were isolated based on tdTomato expression and plated on PLO-laminin coated 384 well plates. Media was collected before treatment (Day 0) and at endpoint (Day 5) and Nluc activity in the media was measured by NanoGlo luciferase assay (Promega). Relative light units (RLU) were read with a microplate reader (ClariOstar, BMG Labtech).

### Cell isolation and culture from human surgical samples

Tissue samples were derived from adult and pediatric patients and prepared for RNA sequencing studies as previously described,^44–46^ with use approved by the research ethics boards of the Montreal Neurological Institute and the Montreal Children’s Hospital. In brief, non-diseased areas of predominantly white matter resected during surgical procedures were processed for cell isolation within 2 h. Samples were dissociated and A2B5^-^ cells were selected with A2B5 antibody conjugated microbeads (Miltenyi Biotec). Cells were treated with IFN-γ for 2 days and RNA was extracted bulk RNA-sequencing using Illumina platform and the NovaSeq 6000 PE100 machine at the Génome Québec Centre.

### Immunocytochemistry

OPCs were cultured on coverslips coated with PDL. Cells were washed with PBS and fixed with 4% PFA for 10 min. Cells were then permeabilized with 0.1% Triton X/PBS for 10 min before blocking with 3% BSA/PBS for 1 hr. Primary antibodies were incubated in 1% BSA/PBS at 4 °C overnight. Secondary antibodies were incubated in 1% BSA/PBS with 4′, 6-diamidino-2-phenylindole (DAPI) for 1 hr at RT. Coverslips were mounted on slides and imaged on Zeiss Axio Observer Z1 with 20x objective. 3x3 tile images from four random fields were imaged per coverslip. Quantification was performed in a blinded manner. Primary antibodies used included: goat-anti-Olig2 (1:250, AF2418, R&D Systems), mouse-anti-CC1 (1:100, OP80, Millipore), rabbit-anti-Olig2 (1:250, AB9610, Millipore), rat-anti-MBP (1:50, MAB386, Millipore), and anti-mouse H2Kb-PE (1:100, 553570, BD Pharmingen). Secondary antibodies used included: donkey-anti-goat Alexa Fluor^®^ 488 (A32814, Invitrogen) and donkey-anti-rabbit Alexa Fluor^®^ 488 (A21206, Invitrogen), donkey-anti-mouse Alexa Fluor^®^ 555 (A31570, Invitrogen), and chicken-anti-rat Alexa Fluor^®^ 647 (A21472, Invitrogen).

### Flow cytometry

#### OPCs

OPCs were detached from the plate with Accutase (Millipore) and centrifuged. Cells were washed with PBS and then blocked with TruStain FcX anti-mouse (BioLegend) in FACs buffer (0.5% BSA/PBS) for 10 min before antibody staining with anti-mouse CD140a-BV605 (1:100, 135916, BioLegend), anti-mouse O4-APC (1:50, 130.118.978, Miltenyi), anti-mouse CD45-APC/Fire750 (1:500, 103154, BioLegend), anti-mouse ACSA2-PE-Vio770, anti-mouse H2Kb-PE (1:400, 553570, BD Pharmingen), and anti-mouse IA IE-BV785 (1:100, 107645, BioLegend). OPCs were stained with antibodies for 20 min in the dark before washing with FACs buffer and centrifuged. Cells were then resuspended in FACs buffer with Propidium Iodide (PI) for flow cytometry with Cytek Aurora 4 laser flow cytometer.

#### CD8^+^ T cells

CD8^+^ T cells were collected in the media and spun down. Cells were washed with PBS and resuspended in Live/Dead aqua (1:1200, L34957, Invitrogen) for 20 min. Cells were then blocked with TruStain FcX anti-mouse (BioLegend) for 10 min before staining cell surface markers including anti-mouse CD8-APC/Fire750 (1:500, 100766, BioLegend), anti-mouse TCR Vβ5.1, 5.2-FITC (1:500, 139514, BioLegend), anti-mouse CD44-BV785 (1:500, 103059, BioLegend), anti-mouse CD25-PE-eFluor610 (1:500, 61-0251-82, Invitrogen), anti-mouse CD69-BV605 (1:500, 104530, BioLegend). Cells were washed with FACs buffer and fixed with Intracellular Fixation Buffer (eBioscience). Cells were then permeabilized with Permeabilization buffer (Cell Signaling Technologies) prior to intracellular antibody staining for 30 min including anti-mouse IFN-γ-APC (1:200, 17-7311-82, Invitrogen), anti-mouse Perforin-PE (1:200, 12-9392-82, Invitrogen), anti-mouse Granzyme B-PE-Cy7 (1:200, 25-8898-82, Invitrogen). Cells were finally washed and resuspended in FACs buffer for flow cytometry on Cytek Aurora 4 laser flow cytometer.

#### Tissue

Animals were perfused with ice-cold HBSS^-/-^ (Gibco) and the corpus callosum was dissected out from the brain. Tissue was minced and dissociated into single cells with collagenase IV (Worthington). Myelin and dead cells were removed with Debris removal solution (Miltenyi) before cells were proceeding to cell staining. Cells were washed with PBS and resuspended in Live/Dead aqua (1:1200, L34957, Invitrogen) for 20 min. Cells were then blocked with TruStain FcX anti-mouse (Biolegend) for 10 min. Cell surface antibodies included: anti-mouse CD45-BV605 (1:200, 103139, BioLegend), anti-mouse CD11b-Pacific Blue (1:1000, 101224, BioLegend), anti-mouse CD3-APC/Fire750 (1:200, 100362, BioLegend), anti-mouse CD4-Spark

NIR 685 (1:200, 100476, BioLegend), anti-mouse CD8-Alexa Fluor 532 (1:200, 58-0081-80, Invitrogen), anti-mouse Vβ11-PerCP-Cy5.5 (1:200, 125912, BioLegend), anti-mouse CD44-BV711 (1:200, 103057, BioLegend), anti-mouse CD62L-PE-Cy7 (1:200, 104428, BioLegend), anti-mouse MHC Class II (I-A/I-E)-BV785 (1:200, 107645, BioLegend), anti-mouse CD86-PerCP (1:200, 105025, BioLegend), anti-mouse NK1.1-APC/Fire810 (1:200, 156519, BioLegend) and anti-mouse CD25-PE-eFluor 610 (1:200, 61-0251-82, Invitrogen). Cells were then permeabilized before staining with intracellular markers including anti-mouse FoxP3-Alexa Fluor 700 (1:200, 126421, BioLegend), anti-mouse IFN-γ-FITC (1:200, 505806, BioLegend), and anti-mouse IL-17A-APC (1:200, 17-7177-81, Invitrogen). Cells were then washed and resuspended in FACs buffer for flow cytometry on Cytek Aurora 4 laser flow cytometer.

### Bulk RNA isolation and qRT-PCR

Total RNA was isolated with TRIzol (Invitrogen) and cDNA was synthesized with High-Capacity RNA-to-cDNA Kit (Invitrogen) according to manufacturer’s instruction. Quantitative real-time PCR was performed in triplicates with Power Track SYBR Green Master Mix (Invitrogen) on the QuantStudio™ 5 Real-Time Fluorescent Quantitative PCR System (Applied Biosystems). The relative expression levels of the genes were calculated using the comparative 2^ΔΔ^ Ct method and normalized to *ActB* expression. Primer sequences are listed in supplemental file S1.

### Bulk RNAseq from mouse and rat OPCs

#### Mouse

Total RNA was isolated with TRIzol (Invitrogen) and RNAseq, library preparation, sequencing, and analysis were performed by NovoGene (Sacramento, CA, USA). Raw fast1 files were processed with fastp software and reads containing adapter, plot-N and low-quality reads were removed. HISAT2 (2.2.1) was used to align cleaned reads to GRCm39 (mm39). featureCounts (v2.0.6) was used to count the number of reads mapped to each gene, and Fragments Per Kilobase of transcript sequence per Million mapped fragments (FPKM) was calculated. Differential expression analysis was done with DESeq2 R package (1.42.0) and P-value was adjusted using Benjamini and Hochberg’s methods. GO and KEGG pathway enrichment analysis was performed with clusterProfiler (4.8.1). Gene Set Enrichment Analysis (GSEA) was performed by ranking genes according to the degree of differential expression. The predefined gene set were then tested to observe enrichment.

#### Rat

Cells were washed and then lysed in RLT Plus lysis buffer. RNA was isolated using the Qiagen RNeasy Plus Mini Kit according to the manufacturer’s recommendations. Isolated RNA was quality controlled, and RNA-sequencing libraries were prepared using the Illumina TruSeq Stranded mRNA kit. Libraries were sequenced at the Johns Hopkins University Single Cell Transcriptomics Core on an Illumina NovaSeq 6000 using a 2 × 75 bp configuration. A total of three biological replicates were used, with each biological replicate consisting of an independent pooled litter. Replicates were collected across two different experimental days.

Transcript quantification from sequencing data was performed by pseudoalignment using Salmon^47^ v1.1.0 against the Rnor_6.0 genome obtained from NCBI,^48^ using selective alignment with the entire genome serving as decoys.^49^ Transcript quantifications were imported into R using tximport v1.14.2,^50^ with ‘countsFromAbundanc’ set to ‘lengthScaledTPM’.

Differential expression analysis was performed using the limma package v3.42.2^51^ following established workflows.^52^ Lowly expressed genes were filtered using the ‘filterByExpr’ function and normalized using ‘calcNormFactors’ with the trimmed mean of M-values (TMM) method. Counts were transformed for linear modeling with precision weights using the ‘voom’ function, and gene-wise linear models were fitted using ‘lmFit’. The design formula was ‘∼ 0 + group + replicatè, where group was defined as baseline, T3 + vehicle, or T3 + cytokine. Moderated t-statistics, F-statistics, and log-odds of differential expression were calculated by empirical Bayes moderation of the standard errors using the ‘eBayes’ function.

### Western blotting

Cell lysates were separated on sodium dodecylsulfate (SDS)-polyacrylamide (7–15%) gels (Bio-Rad) and then transferred on to Immobilion-P membranes (Millipore). Membranes were blocked with either 5% skim milk/TBS-T or 5% BSA/TBS-T before being incubated with primary antibodies at 4 °C overnight. Membranes were washed with TBS-T prior to secondary antibody incubation for 1 hr. Blots were developed using the LiCor Odyssey CLX system with SuperSignal West Femto kit (Pierce). Primary antibodies used are with rabbit anti-SREBP1 (1:1000, PA1-337, Invitrogen) and anti-β-actin-HRP (1:20000, MA5-15739-HRP, Invitrogen). goat anti-rabbit IgG (H+L)-HRP (1:5000, G-21234, Thermo Fisher Scientific) was used for secondary antibody.

### Seahorse extracellular flux assay

Oxygen consumption rate (OCR) was measured using a Seahorse XF96 Extracellular Flux Analyzer (Agilent Technologies). 15,000 OPCs were plated on Seahorse XF96 Cell Culture Microplate (Agilent Technologies) coated with PDL. 24 h after treatment, media was changed to XF assay medium supplemented with 10 mM Glucose, 1 mM Pyruvate, and 2 mM Glutamine and incubated in a CO2-free incubator for 1 h. Vehicle or 40 µM Etomoxir, 2.5 µM Oligomycin, 2 µM FCCP, 0.5 µM Rotenone and 0.5 µM Antimycin A was sequentially injected to the well to measure OCR. Experiments were done in at least triplicates and readings were normalized by cell count.

### Untargeted metabolomics

Metabolomics was performed using two methods:

#### Fig. 1C

Protein-normalized cell homogenate volumes were mixed with extraction solvent consisting of 70% ice-cold methanol containing 0.5% 1N HCl. The extraction solvent was pre-spiked with 12 isotopically labeled internal standards: glucose-D7, glucose-6-phosphate-¹³C6, octanoic acid-D15, lauric acid-D23, glutamic acid-D3, citric acid-D4, butyryl-L-carnitine, lauroyl-L-carnitine-D3 chloride, cytosine-D2, tyrosine-D2, L-tryptophan-D5, and caffeine-¹³C2. Samples were vigorously vortexed for 5 min using a TissueLyser LT (Qiagen, Switzerland) at 50 Hz and then centrifuged at 13,000 rpm for 20 min at 4°C. The supernatants containing internal standards and unknown metabolites were collected and evaporated to complete dryness using a vacuum concentrator (Thermo Scientific “Savant SpeedVac” SPD 120P2, Waltham, MA, USA). The residues were then resuspended in 150 µL of 50% ice cold methanol containing 0.1% formic acid and two additional standards, L-leucenol and sulfinpyrazone (2 ppm each) that were used to track instrument performance and mass accuracy throughout the analyses. Samples were vortexed, centrifuged at 13000 rpm at 4C for 10 min, and supernatants collected for metabolomics profiling.

**Figure 1.**
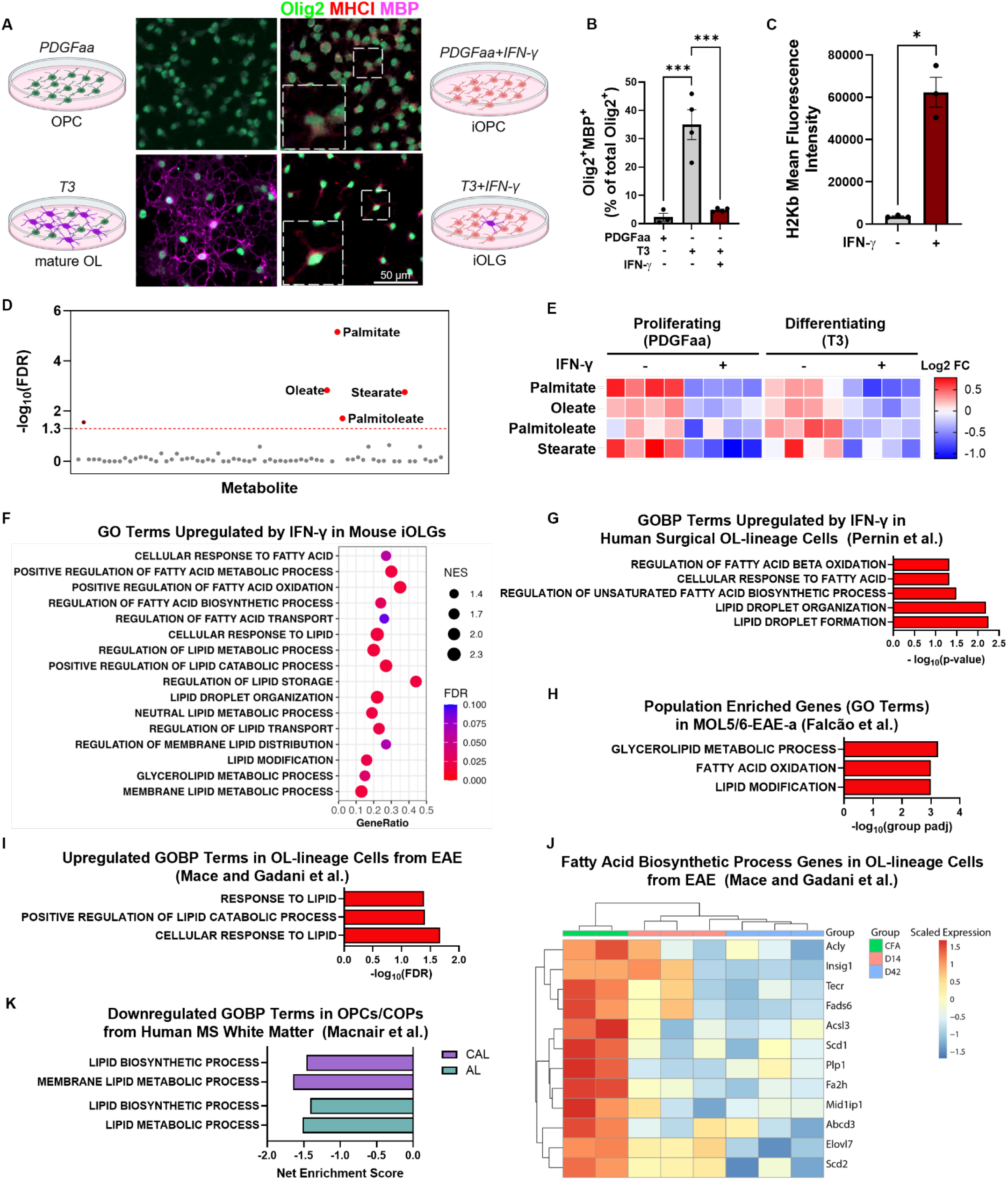
Immune oligodendroglia reprogram lipid metabolism. **(A – C)** Induction of mouse iOLGs with IFN-γ. **(A)** Schematics and representative images of OPCs treated with PDGF-AA, PDGF-AA+IFN-γ, T3, or T3+IFN-γ for 48 h, followed by immunofluorescent staining for Olig2 (marker of OL lineage), MHC class I, and MBP (marker of mature OLs). Inset (large box) represents an enlarged view of the cell shown in the small box. **(B)** Quantification of immunofluorescent staining for MBP. *n* = 4 independent experiments per group, with quantification from four fields per sample. **(C)** Quantification of H2Kb (MHC class I) expression via flow cytometry. *n* = 3 independent experiments performed in triplicate. **(D – E)** Isolated mouse OPCs were treated as in (A) for 24 h, followed by LC-MS metabolite quantification from cell lysates. Data were derived from four biological replicates per condition. **(D)** Shown are metabolites whose concentrations were significantly altered (FDR < 0.05) by IFN-γ treatment, with annotation of free FAs. **(E)** Heatmap of top four metabolites (all free FAs) significantly altered by IFN-γ treatment. **(F)** GSEA showing upregulation of GO terms associated with lipid metabolism following T3+IFN-γ vs T3 treatment of cultured mouse OPCs. *n* = 3 biological replicates per condition. **(G)** GOBP terms upregulated by IFN-γ treatment in surgically derived human OL-lineage cells. Re-analysis from Pernin et al., 2024.^46^ *n* = 3-6 independent samples per group. **(H)** GO terms enriched in EAE-specific subset of early OLs (MOL5/6-EAE-a) from Falcão et al.^17^ **(I)** GOBP terms that were upregulated in OL-lineage cells at peak EAE (day 14) compared to CFA control. Re-analyzed from Mace and Gadani, et al.^55^ **(J)** Heat map showing pseudobulked expression values from OL nuclei, re-analyzed from Mace and Gadani, et al.^55^ Genes shown were significantly downregulated at EAE day 42 (D42) relative to CFA and belong to the GO term “fatty acid biosynthetic process,” which was overrepresented in downregulated genes at both D14 and D42. Each column represents an independent biological replicate. **(K)** Lipid metabolism GO terms significantly downregulated in OPCs+COPs derived from active (AL) and chronic active (CAL) white matter human multiple sclerosis lesions compared to control white matter. Analysis of snRNAseq data from Macnair et al.^21^ Statistical analysis performed via one-way ANOVA with Tukey multiple comparison test (B) and Benjamini-Hochberg FDR correction (D, E), two-tailed Welch’s t-test (C), clusterProfiler adjusted for FDR (F-I), DESeq2 adjusted for FDR (J), or fast gene set enrichment analysis (K). \**P* <0.05, \*\*\**P*<0.001.

UFLC-QTOF-HRMS/MS analysis: Metabolomic profiling was performed using an ultrafast liquid chromatography (Shimadzu, Kyoto, Japan) coupled to a TripleTOF 5600 (AB Sciex, Concord, CA, USA) high resolution mass spectrometer (UFLC-HRMS/MS). The UFLC system consisted of a degasser, a quaternary pump, an autosampler, and a temperature-controlled column compartment. The TripleTOF 5600 mass spectrometer was equipped with DuoSpray Turbo V-dual ESI/APCI source probes. The mass spectrometer used an automated calibrant delivery system that was delivered every 15 sample injections by manufacturer recommended calibrants to maintain the mass accuracy of the instrument below 5 ppm. Ten microliters of each extracted sample were subjected to UFLC-HRMS/MS analysis for metabolomic profiling. Metabolites were chromatographically separated on a Kinetex Pentafluorophenyl (PFP/F5) stationary phase (Phenomenex, Torrance, CA, USA) using a binary gradient mobile phase program (acetonitrile-eluent A and ddH2O-eluent B; both contained 0.1% FA) with the following parameters: 0 – 3 min 100 % eluent B, 3 – 13 min, eluent A increased from 0% to 100%, 13 – 19 min eluent A hold at 100%, 19 – 19.10 eluent B increased to 100% from 0%, and hold for 4 min to allow for column equilibration before the next sample is introduced into the system. The UFLC-separated metabolites were introduced into electrospray ionization chamber for data acquisition in both positive and negative modes over a mass range of 30-900 m/z. Data were acquired and collected in an Information Dependent Acquisition (IDA-HRMS/MS) mode, which attains accurate m/z (mass to charge ratio) of both the pre-cursor and fragments of each metabolite in the samples within a given mass range i.e. 30-900 m/z. The acquired data were processed using Sciex OS - Q1.5 data analysis software (AB SCIEX, Concord, Canada) integrated with the NIST (2017) tandem mass spectral library, and the Sciex accurate MS/MS spectral library 2.0 for peak detection, peak alignment, spectral processing, feature identification based on matching pairs of precursor-fragment ions to the tandem MS databases, and peak quantification. Pooled cell homogenates prepared from all experimental samples were extracted and analyzed using an untargeted LC– HRMS/MS workflow to generate a reliable and reproducible targeted metabolite list. The pooled metabolite extract was analyzed in eight sequential injections on the LC–HRMS/MS system, as described above. To be included in the targeted list, each metabolite was required to be detected in at least 7 of 8 analytical runs with a coefficient of variation (CV) below 20%. Metabolites meeting these reproducibility criteria were then used as a pre-validated targeted list for identification and quantification in the experimental samples.

#### Fig. 3C

OPCs were rapidly washed with 75 mM Ammonium carbonate (Sigma) in HPLC-grade water, adjusted to pH 7.4 with formic acid 3 times. Metabolites were extracted in Cold 40:40:20 Acetonitrile:methanol:water. Extracts were centrifuged at 14000 rpm for 10 min at 4 °C. Supernatants were collected and stored at -80 °C until analysis performed by General Metabolomics (Cambridge, MA). Metabolome profiles of the sample extracts were acquired using flow-injection mass spectrometry. The method described here is adapted from Fuhrer et al 2011. The instrumentation consisted of an Agilent 6550 iFunnel LC-MS Q-TOF mass spectrometer in tandem with an MPS3 autosampler (Gerstel) and an Agilent 1260 Infinity II quaternary pump. The running buffer was 60% isopropanol in water (v/v) buffered with 1 mM ammonium fluoride. Hexakis (1H, 1H, 3H-tetrafluoropropoxy)-phosphazene) (Agilent) and 3-amino-1-propanesulfonic acid (HOT) (Sigma Aldrich). The isocratic flow rate was set to 0.150 mL/min. The instrument was run in 4GHz High Resolution, negative ionization mode. Mass spectra between 50 and 1,000 m/z were collected in profile mode. 5 µL of each sample were injected twice, consecutively, within 0.96 minutes to serve as technical replicates. The pooled study sample was injected periodically throughout the batch. Samples were acquired randomly within plates.

Raw profile data were centroided, merged, and recalibrated using algorithms adapted from Fuhrer et al. 2011.^53^ Putative annotations were generated based on compounds contained in the Human Metabolome Database, KEGG, and ChEBI databases using both accurate mass per charge (tolerance 0.001 m/z) and isotopic correlation patterns.

### Untargeted lipidomics

OPCs were rapidly washed with 75 mM Ammonium carbonate (Sigma) in HPLC-grade water, adjusted to pH 7.4 with formic acid three times. Lipids were extracted from cells in 1:1 (v/v) LC/MS grade isopropanol and methanol. Extracts were centrifuged at 14000 rpm for 10 min at 4 °C. Supernatants were collected and stored at -80 °C until analysis. Metabolite extracts were analyzed by LC-MS/MS by General Metabolomics (Cambridge, MA, USA). 2 μL of each sample was injected and separated by UHPLC using a Nexera UHPLC system (DGU-405 degasser unit, LC40DX3 solvent delivery system, SIL-40CX3 auto sampler, CBM-40 system controller, CTO-40C column 3 oven; Shimadzu). Separation was achieved by reverse-phase liquid chromatography using an Acquity Premier CSH C18 column with VanGuard FIT (1.7 μm, 2.1 mm X 50 mm; 186009463, Waters). Separation was achieved using a 5-minute multi-phase linear gradient of the following buffers: Buffer A) 6:4 acetonitrile: H2O (v/v) 1 mM ammonium acetate, B: 9:1 isopropyl alcohol: acetonitrile (9:1, v/v) with 1 mM ammonium acetate. Samples were ionized using an Optimus Turbo V + Dual TIS ion source (Sciex) and were analyzed using an X500R mass spectrometer (Sciex). Samples were analyzed in positive ionization mode and were acquired using a high-resolution data dependent acquisition method.

LC-MS/MS data processing and ion annotation was performed according to accepted protocols for mass spectrometry data processing and feature annotation.^54^ Briefly, annotation was based on matching of chromatographic retention times, MS1 values from detected features, and the resulting MS2 fragment ions to expected retention time ranges for the indicated lipid classes. Annotation was provided as MS1/MS2 when a cosine similarity score of >0.5 with at least two matched signals was observed. An m/z difference of 5 mDa or 3 ppm was allowed for precursor m/z matched, and fragments were allowed a 5 mDa or 5 ppm m/z difference. For MS1/rt matches, a 3 mDa or 3 ppm m/z tolerance was allowed, and the signal needed to elute within 0.3 min of the expected retention time for that lipid or lipid class. Observed peak heights were normalized to the sum of total annotated feature intensities per sample.

### Adoptive transfer – cuprizone mouse model

8–10-week-old C57BL/6 mice were fed 0.2% cuprizone (bis(cyclo-hexanone) oxaldihydrazone) (Sigma) mixed with powdered, irradiated 18% protein rodent diet (Teklad Global) for 3 weeks, and chow was replaced every 1-2 days. Feed was changed back to normal chow throughout the rest of the experiment. Polarized Th17 cells were injected 1 week after the feed was changed to normal chow.

Spleens and lymph nodes were collected from 2D2^+^ mice and filtered through a 100 µm filter. Cells were pelleted and red blood cells (RBCs) were lysed with RBC lysis buffer (Biolegend). CD4^+^ cells were isolated with CD4^+^ isolation kit (BioLegend) and cocultured with irradiated wild-type splenocytes. Cells were kept in IMDM media (Gibco) added with fetal bovine serum (Gemini Bio-Products), penicillin-streptomycin (Gibco), 2-mercaptoethanol (Gibco), Glutamax (Gibco), and sodium pyruvate (Sigma). To polarize CD4^+^ cells into Th17 cells, 2.5 µg/mL anti-CD3 (BioLegend), 20 µg/mL anti-IL-4 (BioLegend), 20 µg/mL anti-IFN-γ (BioLegend), 30 ng/mL IL-6 (PeproTech), and 3 ng/mL TGFb (Thermo Fisher) were added to the media for 48 hrs. IL-23 (BioLegend) was added to maintain Th17 polarity along with fresh media for another 24 hrs. Cells were transferred to a new plate with fresh media and IL-23 to rest for 48 hrs. Cells were restimulated on plates coated with anti-CD3 and anti-CD28 (BioLegend) for 40 hrs before harvest. Cells were resuspended in PBS and counted. A final number of 10 million viable cells/250 µl were adoptively-transferred via IP.

### Immunohistochemistry

Brain sections were sliced at 30 µm sections on Leica CM1850 cryostat (Leica Biosystems) and washed with PBS. Sections were incubated in Liberate Antibody Binding Solution (Polysciences) for 10 min. After two washes in PBS, sections were permeabilized in 0.5% Triton-X 100/PBS for 10 min Sections were then blocked in 10% blocking solution (10% normal donkey serum, 3% Triton-X 100 in PBS) for 1 hr and incubated with primary antibodies at 4 °C overnight. Sections were washed three times with PBS and incubated with secondary antibodies and DAPI in 10% blocking solution for 3 hr at RT. After three washes with PBS, sections were mounted on slides and imaged with Zeiss LSM 900 confocal. Quantification was performed in a blinded manner. Primary antibodies used included: goat-anti-Olig2 (1:250, AF2418, R&D Systems) and rabbit-anti-ASPA (1:300, PA5-29180, Invitrogen). Secondary antibodies used included: donkey-anti-goat Alexa Fluor^®^ 647 (A21447, Invitrogen) and donkey-anti-rabbit Alexa Fluor^®^ 555 (A32794, Invitrogen). Myelin was stained with FluoroMyelin™ Green (1:50, F34651, Invitrogen).

### Re-analysis of Mace and Gadani et al. EAE spinal cord snRNAseq data

The previously published dataset^55^ (Gene Expression Omnibus accession GSE282120) was reanalyzed to focus on oligodendrocytes for this study. Raw UMI counts generated by Cell Ranger (v7.2.0) from wild-type CFA (*n* = 2), day 14 EAE (D14; *n* = 3), and day 42 EAE (D42; *n* = 3) samples were imported into R (v4.5.1), subsetted, and analyzed using Seurat^56^ (v5.3.0). Likely doublets were identified using scDblFinder^57^ (v1.22.0).

Using Seurat, per-nucleus library size was normalized by dividing feature counts by total counts, multiplying by a scale factor of 10,000, adding 1, and natural log-transforming. Highly variable features were identified using Seurat’s default variance-stabilizing transformation method and were subsequently scaled. Dimensionality reduction was performed by principal component analysis on the 2,000 scaled variable features. The first 28 principal components were used to construct a nearest-neighbor graph with k = 20, followed by Louvain clustering at a resolution of 0.2.

Cell types were programmatically identified using SingleR^58^ (v2.10.0) with the mouse mRNA-seq reference^59^ provided by the celldex package^58^ (1.18.0). Clusters identified as containing a high proportion of oligodendrocytes were manually confirmed by expression of oligodendrocyte-associated transcripts, including *Opalin*, *Klk6*, *Mbp*, *Mobp*, *Mag*, *Mog*, *Plp1*, and *Olig2*.

Established outlier detection metrics were then used to identify poor-quality nuclei on a per-sample basis within oligodendroglial lineage nuclei.^60^ Nuclei were considered low quality if two or more of the following criteria were met: library size, defined by either UMI counts or detected features, greater than 2.5 median absolute deviations (MADs) above the median or 5 MADs below the median per sample on a log10 scale; percentage of reads mapping to the top 20% of expressed genes greater than 5 MADs above the median; deviation greater than 5 MADs above or below the predicted feature-count linear regression; percentage of reads mapping to the mitochondrial genome greater than 2.5 MADs above the median; or greater than 1% of reads mapping to the mitochondrial genome.

Oligodendroglial lineage nuclei were then analyzed as described above for the full dataset, except that only the first 15 principal components were used to construct the nearest-neighbor graph. Clusters enriched for low-quality nuclei, doublets, or nuclei expressing genes not associated with oligodendroglia were removed, along with any remaining low-quality nuclei and predicted doublets. The violin plot in Supplementary Fig. 7B was generated in Seurat using these purified log-normalized nuclei.

For differential expression and pathway enrichment analyses, oligodendrocyte nuclei were aggregated into individual pseudobulk samples, with one pseudobulk sample per biological replicate. All eight biological replicates contained more than 600 oligodendrocyte nuclei prior to aggregation. Pseudobulk samples were analyzed using edgeR’s quasi-likelihood method^61^ (v4.6.3), as implemented in the scran function *pseudoBulkDGE*^62^ (v1.36.0), and separately using DESeq2^63^ (v1.48.2). The design formula was*∼0 + group*, where group was one of CFA, D14, and D42. Three contrasts were tested: D14 versus CFA, D42 versus CFA, and D42 versus D14.

To identify pathways enriched at the two EAE time points, we used the competitive gene set test CAMERA from the limma package^64^ (v3.64.3) on the edgeR quasi-likelihood results. Gene set testing was performed against Biological Process gene sets from the Gene Ontology database,^65,66^ which were retrieved from the Molecular Signatures Database (MSigDB) (MSigDB 2026.1.Mm).^67^ In DESeq2, genes with an adjusted *P* -value less than 0.05 were considered differentially expressed.

The heatmap in Fig. 1H was generated using the pheatmap package (v1.0.13) from normalized and variance-stabilized counts, obtained using the *vst* function in DESeq2. The heatmap included genes that were downregulated in D42 compared with CFA and associated with the GO Biological Process term “Fatty acid biosynthetic process” GO:0006633.

### Analysis of MacNair et al. post-mortem human multiple sclerosis snRNAseq data

The cleaned and annotated count data, as well as the original authors’ published analysis code from MacNair et al.,^21^ were retrieved from Zenodo (https://doi.org/10.5281/zenodo.8338963). After loading the data into R and subsetting to include only OPC-COP and oligodendrocyte nuclei, the data were analyzed using the previously published code with only one modification: the edgeR function *getNormLibSizes* was used to replace the now-deprecated function *effectiveLibSizes*. The published analysis script uses glmmTMB^68^ (v1.1.14) to identify differentially expressed genes between lesion types for broad cell types and fgsea^69^ (v1.34.2) to perform gene set enrichment analysis on the differentially expressed genes.

### Cell proliferation and viability assays

15,000 OPCs were plated on PDL coated 96 well black plate (Corning). Cells were stained with Incucyte® Nuclight Rapid Red Dye (1:1500) and Incucyte® Cytotox Dye (1:15000) for all cells and dead cells, respectively, at the time of treatment. Cells were imaged with Sartorius Incucyte live cell imager every 6 h. After 72 h, Incucyte software was used to analyze the number of Nuclight Red positive cells and Cytotox positive cells. Viable cell numbers were calculated by the number of Nuclight Red positive cells and Cytotox negative cells. Viability (%) was calculated by the ratio of Nuclight Red positive cells and Cytotox negative cells to all Nuclight Red positive cells.

### Statistical analyses

For RNAseq analyses, differential expression was evaluated using the DESeq2 R package (1.42.0) and P-value was adjusted using Benjamini and Hochberg’s methods. KEGG and GO pathway enrichment analysis was performed with clusterProfiler (4.8.1) and terms with FDR lower than 0.05 were considered significantly enriched. For all other studies, statistical analyses were performed using Prism 11.0 (GraphPad Software Inc.). *P*<0.05 was considered statistically significant. The statistical tests used are detailed in the figure legends and included two-tailed Welch’s t-test when comparing two groups or one-way or two-way ANOVA followed by Dunnett’s or Tukey’s post hoc test when comparing more than two groups. Sample size and number of replicates are also described in the figure legends.

## Results

### iOLGs reprogram lipid metabolism

To understand the metabolic changes induced by inflammatory cytokine exposure in OPCs, we first performed untargeted metabolomics in primary cultured mouse OPCs that were kept in proliferating media (including platelet-derived growth factor-aa, or PDGF-AA) or differentiating media (including triiodothyronine, or T3) with or without IFN-γ treatment. IFN-γ is a potent inducer of iOLGs, and consistent with prior studies^11,16^ we found that IFN-γ induced expression of MHC class I under both proliferative and differentiating conditions while preventing differentiation, as measured by CC1 and myelin basic protein (MBP) expression (Fig. 1A-C and Supplementary Fig. 1). Metabolomic analysis of cell lysates revealed that the free FAs palmitate, oleate, palmitoleate, and stearate were the most significantly changed metabolites in response to IFN-γ (Fig. 1D). All four FAs were significantly reduced by IFN-γ treatment in OPCs under both proliferative and differentiating conditions (Fig. 1E).

To gain further insight into the metabolic changes induced by IFN-γ, we performed RNA sequencing (RNAseq) from mouse OPCs treated with or without IFN-γ for 24 h in the presence of T3. Consistent with our metabolomics analysis, many transcriptional programs associated with lipid and FA metabolism were significantly upregulated by IFN-γ, including Gene Ontology (GO) terms linked to regulation of the biosynthesis, storage, and intracellular localization of lipids, along with positive regulation of FA β-oxidation (FAO) and lipid catabolism (Fig. 1F). A second, independent RNAseq analysis performed from rat OPCs produced strikingly similar results, with IFN-γ leading to upregulation of pathways associated with FAO and lipid catabolism along with broad changes in lipid transport and localization (Supplementary Fig. 2).

To determine whether the effects of IFN-γ on lipid and FA metabolism are conserved in human iOLGs, we analyzed published RNAseq data from cultured OL-lineage cells isolated from pathologically normal tissue within human surgical specimens.^44–46^ Similar to rodent cells, exposure of surgically derived human OLs to IFN-γ produced broad changes in transcriptional programs associated with lipid metabolism, including those linked to FAO, FA synthesis, and lipid storage and organization (Fig. 1G).

We next investigated the relevance of these findings to disease pathology by examining whether similar changes in lipid metabolic programs are observed in mouse models of inflammatory demyelination and in human multiple sclerosis tissue. To do so, we examined published RNAseq datasets. A single-cell RNA sequencing (scRNAseq) analysis performed from the spinal cords of mice subjected to the experimental autoimmune encephalomyelitis (EAE) mouse model of multiple sclerosis (Falcão, et al.)^17^ identified a disease-specific population of early OL-lineage cells (termed MOL5/6-EAE-a) defined by transcriptional signatures associated with lipid metabolism, including FAO (Fig. 1H). Analysis of a second, independent single-nucleus RNA sequencing (snRNAseq) dataset from mouse EAE (Mace and Gadani, et al.)^55^ similarly showed changes in lipid metabolic programs in disease-associated OL populations, with upregulation of terms associated with lipid response and lipid catabolism (Fig. 1I) along with downregulation of FA biosynthesis genes (Fig. 1J). Lastly, we analyzed a snRNAseq dataset derived from human multiple sclerosis lesions (Macnair, et al.).^21^ Gene programs associated with lipid metabolism were altered in OPCs and committed oligodendrocyte precursors (COPs) from both active and chronic active lesions when compared to control white matter, including downregulation of lipid biosynthesis genes (Fig. 1K).

Together, these findings demonstrate that inflammatory signals produce a broad reprogramming of lipid metabolism in OL-lineage cells, both *in vitro* (in rodent and human cells) as well as in mouse models and human multiple sclerosis tissue. These changes are characterized by a decrease in intracellular free FA levels and downregulation of lipid biosynthesis gene programs concomitant with upregulation of transcriptional programs associated with lipid utilization, such as FAO.

### Fatty acid metabolism modulates OPC responses to IFN-γ

Given that transcriptional programs associated with FAO were increased in iOLGs both *in vitro* and *in vivo*, we first postulated that upregulation of FAO might contribute to the decrease in free FAs we observed in cultured iOLGs. To functionally measure FAO in iOLGs, we performed Seahorse extracellular flux analysis in cultured mouse OPCs exposed to IFN-γ under differentiating conditions (Fig. 2A-C). Although total maximal respiration decreased slightly following IFN-γ treatment (Fig. 2B), the contribution of FAO to respiratory capacity increased, indicating increased dependence on FAO (Fig. 2C).

**Figure 2.**
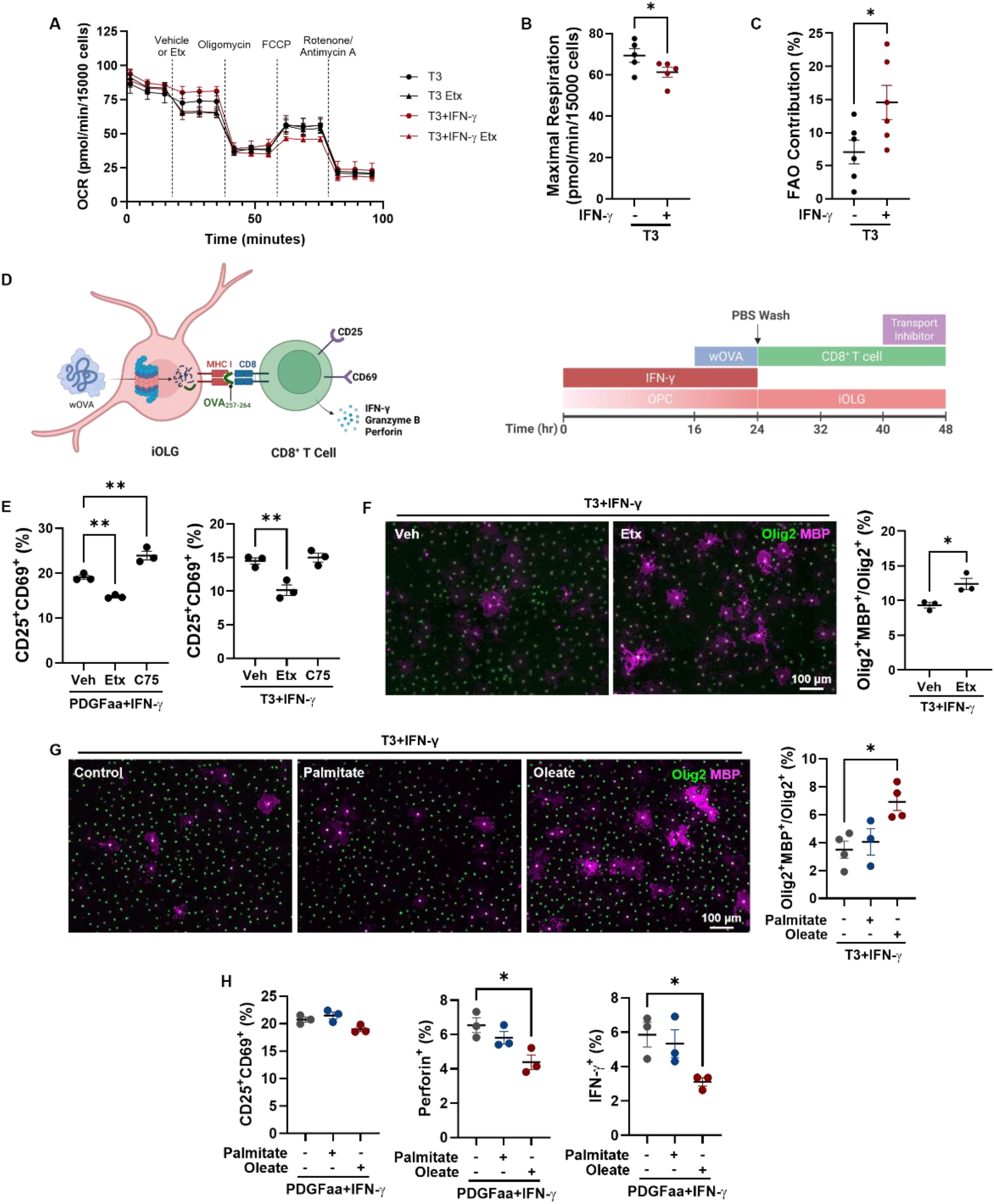
Fatty acid metabolism modulates OPC responses to IFN-γ. **(A – C)** Cultured mouse OPCs were treated with T3 or T3+IFN-γ for 24 h, followed by Seahorse extracellular flux assay to measure oxygen consumption rate (OCR). Etomoxir (Etx) was added to measure the contribution of FAO. *n* = 5-6 independent experiments, each including four technical replicates per group. **(A)** Representative OCR graph from a single experiment. **(B)** Maximal respiration normalized by cell number. **(C)** FAO contribution to total respiration. **(D)** Schematic of iOLG – OT-I CD8^+^ T cell co-culture. Mouse OPCs were stimulated with IFN-γ to induce iOLGs for 16 h. Whole ovalbumin (wOVA) was then added as antigen for iOLGs to phagocytose and process for cross-presentation. Cultures were washed, and CD8^+^ T cells isolated from OT-I mice were added for 24 h before flow cytometric analysis. **(E)** iOLGs were induced with IFN-γ in the presence of vehicle, Etx (40 µM), or C75 (20 µM) prior to wOVA supplementation, PBS wash, and OT-I CD8^+^ T cell co-culture. CD8^+^ T cell activation was then determined by CD25^+^CD69^+^ expression gated on CD8^+^Vβ5^+^ cells via flow cytometry. *n* = 3 independent experiments performed in triplicate. **(F)** Mouse OPCs were cultured in T3+IFN-γ ± Etx (40 µM) for 72 h, followed by immunofluorescent staining for Olig2 and MBP. Representative images (*left*) and quantification of MBP^+^Olig2^+^ cells (*right*). *n* = 3 independent experiments. **(G)** Mouse OPCs were cultured in T3+IFN-γ with 50 µM bovine serum albumin (BSA)-conjugated-palmitate, -oleate, or BSA-alone control for 48 h, followed by immunofluorescent staining for Olig2 and MBP. Representative images (*left*) and quantification of MBP^+^ Olig2^+^ cells (*right*). *n* = 4 independent experiments. **(H)** iOLGs were induced with IFN-γ in the presence of BSA-conjugated-palmitate, -oleate, or BSA-control before OT-I CD8^+^ T cell co-culture. CD8^+^ T cell activation was measured by CD25^+^ CD69^+^, Perforin^+^, and IFN-γ^+^ expression via flow cytometry. *n* = 3 independent experiments performed in triplicate. \**P* <0.05, \*\**P*<0.01 by two-tailed Welch’s t-test (B, C, F) or one-way ANOVA with Dunnett multiple comparison test (E, G, H). For immunofluorescence studies (F, G), each experiment consisted of independent biological samples with quantification from four fields per sample. Graphs shown as mean ± SEM.

We next asked whether the increased dependence on FAO represented a metabolic vulnerability that could be targeted to modulate the functional effects of IFN-γ on OPCs. As noted above, iOLGs express MHC class I and are capable of phagocytosing, processing, and cross-presenting antigen to activate CD8^+^ T cells *in vitro.*^16^ We assessed this immunologic function using a co-culture assay, in which IFN-γ-induced iOLGs were supplemented with whole ovalbumin (wOVA), which requires phagocytosis and processing prior to antigen presentation, followed by co-culture with CD8^+^ T cells isolated from OT-I mice (Fig. 2D). OT-I mice express transgenic T cell receptors designed to recognize OVA257-264 peptide presented by the MHC I molecule, thereby producing MHC class I-restricted, OVA-specific CD8^+^ T cells.^70^ We then measured CD8^+^ T cell activation via flow cytometry (Supplementary Fig. 3A and B). Treatment of IFN-γ-induced iOLGs with etomoxir (40 µM), an FAO inhibitor that targets the rate-limiting enzyme carnitine palmitoyltransferase 1a (Cpt1a), attenuated activation of co-cultured CD8^+^ T cells as assessed by CD25 and CD69 expression (Fig. 2E and Supplementary Fig. 3C) without affecting iOLG cell viability (Supplementary Fig. 3D). Conversely, CD8^+^ T cell activation increased when iOLGs were treated with the FA synthesis inhibitor and FAO activator C75.^71^ Moreover, inhibition of FAO with etomoxir attenuated IFN-γ-associated OL differentiation blockade, as assessed by MBP expression (Fig. 2F).

The above findings suggest that increased FAO plays a direct role in mediating the functional effects of IFN-γ on OPCs, including induction of immune-like functions and inhibition of OL differentiation. Conversely, we wondered whether the observed depletion of free FAs might also have functional relevance, given that these FAs represent critical building blocks for myelin membrane synthesis. We therefore tested whether supplementation of the depleted FAs can reverse the functional effects of IFN-γ. Intriguingly, supplementation of oleate but not palmitate consistently enhanced OL differentiation, partially overcoming IFN-γ-induced differentiation blockade (Fig. 2G). We confirmed that supplementation of FAs did not change cell viability (Supplementary Fig. 3E). Similarly, supplementation of IFN-γ-treated OPCs with oleate, but not palmitate, attenuated inflammatory activation of co-cultured CD8^+^ T cells (Fig. 2H). While oleate supplementation of iOLGs did not significantly impact induction of CD25^+^ CD69^+^ expression, it led to significantly decreased production of perforin and IFN-γ in co-cultured CD8^+^ T cells.

### LXR activation reverses effects of IFN-γ on OPC lipid metabolism and overcomes differentiation blockade

Although FAO and FA depletion were found to be targetable biological processes for modulating OPC responses to inflammation, the effects of FAO inhibition and oleate supplementation were modest. Moreover, our transcriptomic analyses from cultured OPCs, mouse EAE, and human multiple sclerosis tissue suggested broad inflammation-induced changes in lipid metabolism, extending beyond FAO and FA depletion. These observations prompted us to move from targeting individual metabolic pathways to identifying key upstream transcriptional regulators of iOLG lipid metabolism that might serve as central nodes for therapeutic modulation.

The LXRs are a family of transcription factors consisting of two isotypes, LXRα (encoded by *NR1H3*) and LXRβ (encoded by *NR1H2*). They serve as master regulators of lipid metabolism, influencing FA metabolism (including synthesis, elongation, desaturation, and incorporation into phospholipids) as well as cholesterol handling.^72^ Through effects on lipid metabolism, the LXRs regulate the functions of varied immune subsets, including both myeloid and lymphoid populations, with activation of the LXRs generally producing anti-inflammatory effects.^72–74^ Moreover, the LXRs, and in particular LXRβ, are expressed in OPCs and support mOL differentiation during development and under non-inflammatory conditions, likely through enhancement of lipid and cholesterol synthesis required for myelin membrane formation.^75,76^ Given these functions and their role in promoting lipogenesis, which might oppose the switch toward lipid catabolism we observed with IFN-γ exposure, we investigated the impact of LXRs on lipid metabolism and functional responses in iOLGs. To do so, we utilized the well characterized LXR agonist GW3965, which potently and selectively activates LXR transcriptional activity both *in vitro* and *in vivo.*^77–79^ We cultured primary mouse OPCs under differentiating conditions in the presence of IFN-γ ± GW3965. We confirmed that treatment with GW3965 (20 µM) robustly increased expression of LXR target genes (Supplementary Fig. 4A) without impacting cell proliferation or viability (Supplementary Fig. 4B and C).

To interrogate the effects of LXR activation, we first performed RNAseq from IFN-γ-induced mouse iOLGs ± GW3965. Kyoto Encyclopedia of Genes and Genomes (KEGG) enrichment analysis found that GW3965 upregulated transcriptional programs associated with FA metabolism and lipid synthesis, including terms “Fatty acid metabolism,” “Biosynthesis of unsaturated fatty acids,” and “Fatty acid biosynthesis” (Fig. 3A), consistent with a switch toward lipogenesis. Many key genes involved in FA synthesis were increased (Fig. 3B), including *Fasn* (rate-limiting enzyme in FA synthesis) and *Srebf1* (transcription factor promoting FA synthesis). We further confirmed this increase with qPCR (Supplementary Fig. 4D) and at the protein level (Supplementary Fig. 4E). LXR activation also significantly increased expression of *Scd1* and *Scd2* (Fig. 3B), rate-limiting enzymes in the generation of monounsaturated FAs such as oleic acid, the supplementation of which mitigated the effects of IFN-γ on iOLG immune functions and mOL differentiation (Fig. 2G-H). These genes showed decreased expression in OL-lineage cells derived from EAE mice (Fig. 1J and Supplementary Fig. 4F), which might contribute to the depletion of oleic acid and other unsaturated FAs caused by inflammatory stimuli. Strikingly, and consistent with our RNAseq data, metabolomics analysis showed that LXR activation with GW3965 prevented the depletion of free FAs induced by IFN-γ (Fig. 3C). These findings demonstrate that pharmacologic activation of LXR counteracts the metabolic effects of IFN-γ, stimulating FA synthesis and increasing free FA levels.

**Figure 3.**
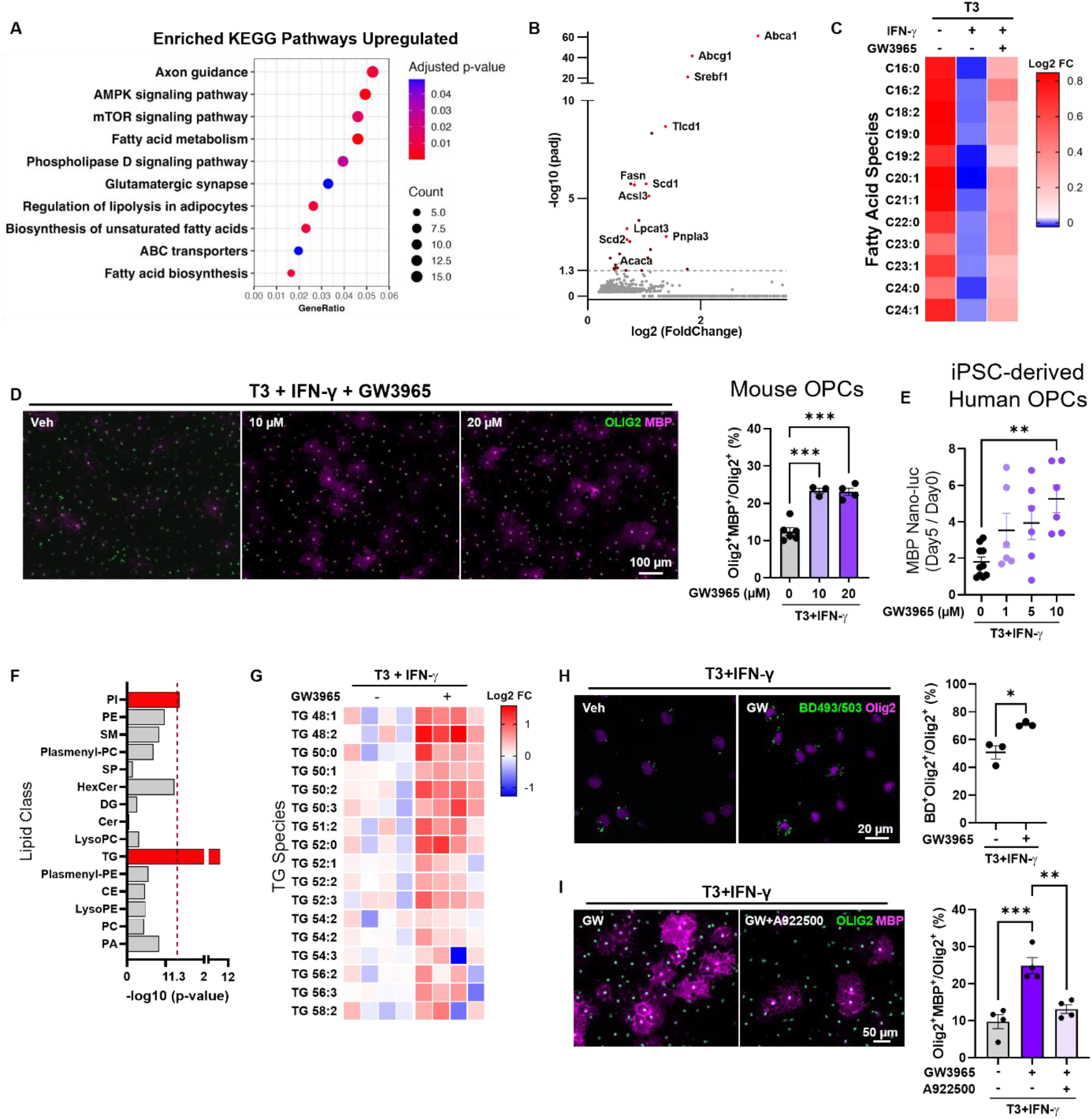
LXR activation reverses effects of IFN-γ on OPC lipid metabolism and overcomes differentiation blockade. **(A – B)** RNAseq analysis of IFN-γ-induced mouse iOLGs treated with the LXR agonist GW3965 versus vehicle for 24 h. *n =* 4 biological replicates. **(A)** KEGG pathways upregulated with GW3965 treatment. **(B)** Volcano plot of genes upregulated with GW3965 treatment. **(C)** Heat map of fatty acid species measured by metabolomics analysis of mouse OPCs treated with T3, T3+IFN-γ, or T3+IFN-γ+GW3965 for 24 h. *n =* 4 biological replicates. **(D)** Mouse OPCs were cultured in T3+IFN-γ with the indicated concentrations of GW3965 for 72 h followed by immunofluorescent staining for Olig2 and MBP. Representative images (*left*) and quantification of MBP^+^Olig2^+^ cells (*right*). *n =* 3 or 4 independent experiments per group. **(E)** hPSC-derived human OPCs expressing an *MBP*-driven luciferase reporter were treated with T3+IFN-γ plus vehicle or the indicated dose of GW3965 for 5 days. MBP expression was quantified as the ratio of Nano-luc signal at Day 5 versus Day 0 of treatment. *n =* 6 technical replicates. **(F)** Lipid class enrichment and **(G)** heat map of triglyceride (TG) species in IFN-γ-induced iOLGs treated with GW3965 compared to vehicle for 24 h. *n =* 4 biological replicates. **(H)** Mouse OPCs were cultured in T3+IFN-γ with vehicle or GW3965 for 24 h and then stained for Olig2 and neutral lipid dye BODIPY (BD) 493/503. Representative images (*left*) and quantification of BODIPY positive OPCs (*right*). *n =* 3 independent experiments. **(I)** iOLGs (T3+IFN-γ) were treated with GW3965 and/or the DGAT1 inhibitor A922500 (20 µM) for 72 h and MBP^+^Olig2^+^ cells were quantified by immunofluorescence. *n =* 4 independent experiments. \**P* <0.05, \*\**P*<0.01, \*\*\**P*<0.001. Statistics performed using clusterProfiler (A) or DESeq2 (B) adjusted for FDR, one-way ANOVA with Dunnett (D, E) or Tukey (I) multiple comparison test, Fisher’s exact test (F), or two-tailed Welch’s t-test (H). For immunofluorescence studies (D, H, I), each experiment consisted of independent biological samples with quantification from four fields per sample. Unless otherwise indicated, GW3965 was used at 20 µM. Graphs shown as mean ± SEM.

We next investigated the functional consequences of modulating lipid metabolism via LXR activation in iOLGs. We first examined the effect of LXR activation with GW3965 on IFN-γ-induced OPC differentiation blockade, finding that drug treatment led to significantly increased maturation of mouse OPCs as measured by MBP expression (Fig. 3D). We then validated this finding using a second, structurally distinct LXR agonist, T091317 (Supplementary Fig. 4G). To determine the relevance of LXR signaling in human cells, we utilized human pluripotent stem cell (hPSC)-derived OPCs that had been engineered to express a luciferase reporter under control of the *MBP* gene promoter.^42^ Treatment with GW3965 in the presence of IFN-γ significantly increased *MBP* reporter expression (Fig. 3E), demonstrating that the reversal of IFN-γ-induced differentiation blockade is conserved in human cells.

We further investigated the impact of LXR activation on lipid metabolism by performing lipidomics from induced mouse iOLGs in culture. Class enrichment analysis revealed that triglycerides were significantly increased following LXR activation with GW3965 (Fig. 3F and G). Triglycerides are a primary component of neutral lipid droplets, which serve as intracellular buffers regulating bioenergetics and lipid availability while limiting lipotoxic stress.^80^ Consistent with our lipidomics findings, fluorescent staining with the neutral lipid dye BODIPY 493/503^81^ showed increased accumulation of neutral lipids within lipid droplet-like structures following GW3965 treatment (Fig. 3H). Inhibition of triglyceride synthesis with A922500, a pharmacologic inhibitor of diacylglycerol O-acyltransferase 1 (DGAT1),^82,83^ attenuated the effect of GW3965 on differentiation (Fig. 3I), indicating that LXR-induced lipid buffering via triglyceride formation contributes to its functional benefit in overcoming differentiation blockade.

### LXR activation suppresses iOLG immune functions

In addition to preventing differentiation blockade, we wondered whether reprogramming of lipid metabolism via LXR might reciprocally limit iOLG immune-like functions. KEGG pathway analysis following GW3965 treatment in IFN-γ-induced iOLGs demonstrated decreased expression of transcriptional programs associated with inflammatory terms such as “Allograft rejection” and “Graft-versus-host disease,” as well as pathways related to antigen cross-presentation such as “Lysosome,” “Phagosome,” and “Antigen processing and presentation” (Fig. 4A). Genes encoding the lysosomal proteases Cathepsin A (*Ctsa*) and Cathepsin L (*Ctsl*), which play key roles in antigen processing,^84,85^ were among the most highly downregulated genes following GW3965 treatment (Fig. 4B).

**Figure 4.**
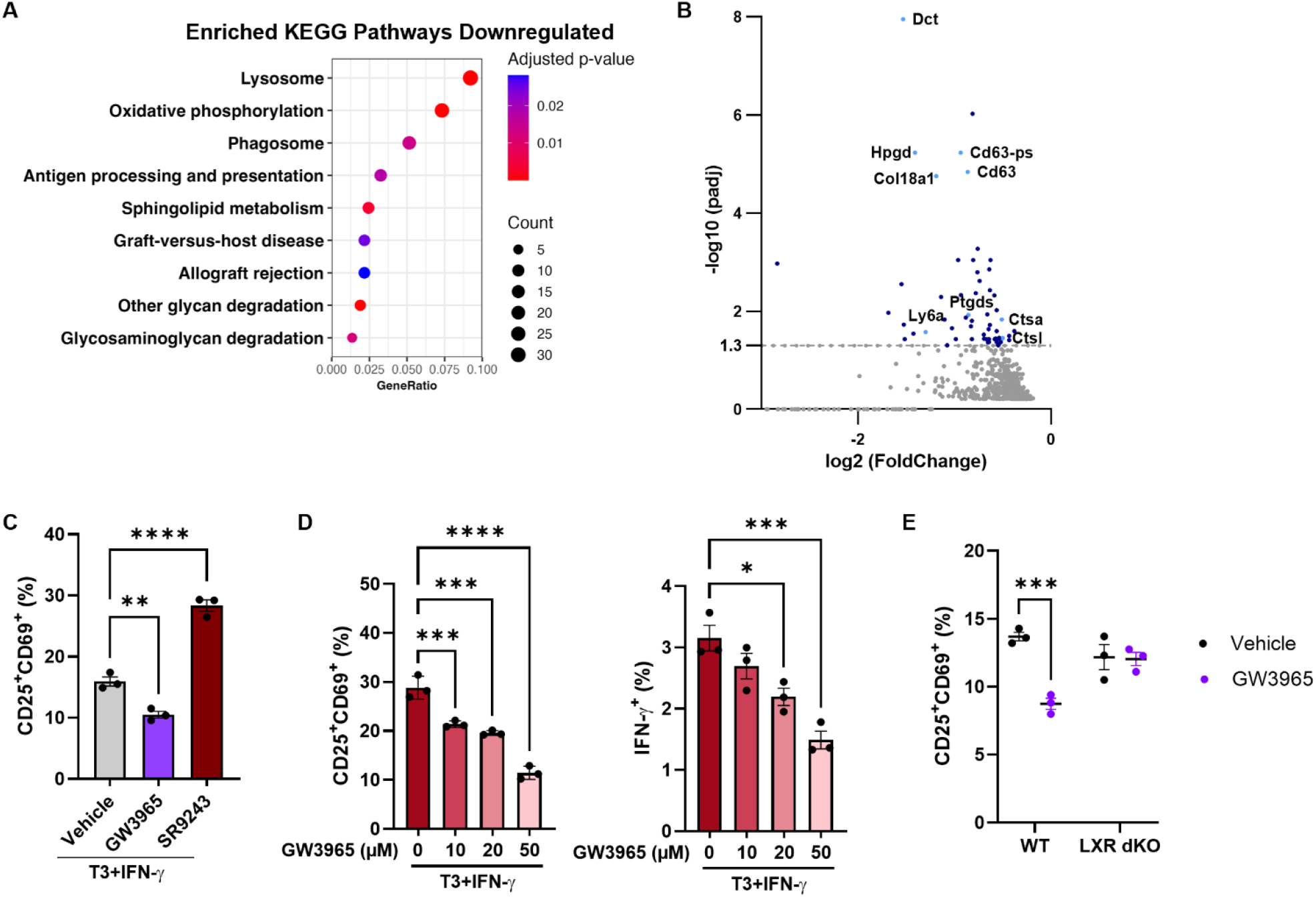
LXR activation suppresses iOLG immune functions. **(A)** KEGG pathways downregulated in mouse iOLGs treated with GW3965 versus vehicle for 24 h. **(B)** Volcano plot of genes whose expression was downregulated in mouse iOLGs treated with GW3965 versus vehicle. *n =* 4 biological samples for both (A) and (B). **(C – E)** iOLG – CD8^+^ T cell co-culture assays. Mouse iOLGs were induced with IFN-γ in the presence of the indicated drugs prior to wOVA supplementation, PBS wash, and co-culture with OT-I CD8^+^ T cells for 24 h. CD8^+^ T cell activation was then measured via flow cytometry. *n =* 3 independent experiments performed in triplicate. **(C)** iOLGs were treated with vehicle, the LXR agonist GW3965, or the LXR inverse agonist SR9243 (10 µM). **(D)** Dose-dependent effect of GW3965 on iOLG-mediated CD8^+^ T cell activation, as measured by CD25^+^CD69^+^ and IFN-γ^+^ expression. **(E)** Wild-type or LXR dKO iOLGs were treated with vehicle or GW3965 prior to co-culture with OT-I CD8^+^ T cells. \**P* <0.05, \*\**P*<0.01, \*\*\**P*<0.001. Statistics performed using clusterProfiler (A) or DESeq2 (B) adjusted for FDR, one-way ANOVA with Dunnett multiple comparison test (C, D), or two-way ANOVA (E). Unless otherwise indicated, GW3965 was used at 20 µM. Graphs shown as mean ± SEM.

To directly examine the effects of LXR activation on iOLG immune functions, we again assayed the ability of wOVA-loaded iOLGs to activate co-cultured OT-I CD8^+^ T cells. Consistent with our transcriptomics analysis, treatment of iOLGs with the LXR agonist GW3965 attenuated their capacity to activate CD8^+^ T cells (Fig. 4C). Conversely, the LXR inverse agonist SR9243 (which inhibits LXR transcriptional activity) had the opposite effect, augmenting cross-presentation and activation of CD8^+^ T cells. The effect of GW3965 on iOLGs was dose-dependent, decreasing CD8^+^ T cell activation markers (CD25 and CD69 expression) and IFN-γ production (Fig. 4D and Supplementary Fig. 5). The suppressive effect of GW3965 was abolished in LXRα and LXRβ double knockout (LXR dKO) iOLGs (Fig. 4E and Supplementary Fig. 6), confirming that drug effects were mediated by LXR.

Taken together, our results demonstrate that LXR activation reprograms lipid metabolism, thereby redirecting iOLGs from an immune-like to a remyelinating phenotype.

### LXR expression in OL-lineage cells under inflammatory conditions

Since LXR activation attenuated the effects of IFN-γ on OPC lipid metabolism and function, we wondered whether LXR isoform expression changed in response to inflammatory stimuli, hypothesizing that LXR expression might decrease in iOLGs. Unexpectedly, expression of both *Nr1h2* (LXRβ) and *Nr1h3* (LXRα) increased in mouse OPCs in response to IFN-γ exposure (Supplementary Fig. 7A). Consistent with this finding, analysis of scRNAseq data (Mace and Gadani, et al.)^55^ found that *Nr1h2* expression was increased in OL-lineage cells from mouse spinal cord at peak EAE compared to controls (Supplementary Fig. 7B), suggesting a similar response *in vivo*. We again analyzed snRNAseq data from human multiple sclerosis tissue (Macnair, et al.)^21^ and similarly found increased expression of both *NR1H2* and *NR1H3* in OLs from active and chronic active lesions (Supplementary Fig. 7C). The discordance between LXR expression and the metabolic and functional consequences of LXR activation suggests that increased expression might be a compensatory response to metabolic reprogramming induced by inflammatory stimuli. Such a compensatory increase in LXR isoform expression has been reported previously in both peripheral and CNS myeloid cells.^72,86–88^

### The LXR agonist GW3965 enhances OPC differentiation and augments remyelination in an adoptive transfer-cuprizone mouse model

To determine whether LXR activation represents a viable strategy for overcoming the inhibitory effects of inflammation on remyelination *in vivo*, we used the adoptive transfer-cuprizone (AT-Cup) mouse model, which combines cuprizone-induced demyelination with adoptive transfer of myelin-targeted Th17 CD4^+^ T cells from 2D2^+^ TCR mice.^16,19,89^ Adoptively transferred CD4^+^ T cells infiltrate the demyelinated corpus callosum and induce a broad CNS immune response (including CD8^+^ T cells), delaying spontaneous remyelination and thereby modeling remyelination in an inflammatory environment. Importantly, iOLGs are abundantly generated in the AT-Cup model,^16,19^ which has advantages for studying immune-mediated inhibition of remyelination over EAE (in which remyelination is patchy and limited by extensive axonal injury) and traditional cuprizone or lysolethicin-mediated demyelination (in which spontaneous remyelination occurs efficiently with a limited inflammatory response). In our experiments, mice were fed 0.2% cuprizone for three weeks to induce demyelination, returned to normal feed for one week, and then adoptively transferred with 2D2^+^ Th17 cells (Fig. 5A). As previously reported,^89^ we validated that adoptive transfer delayed remyelination while inducing a robust immune response within the corpus callosum (Fig. 5B).

**Figure 5.**
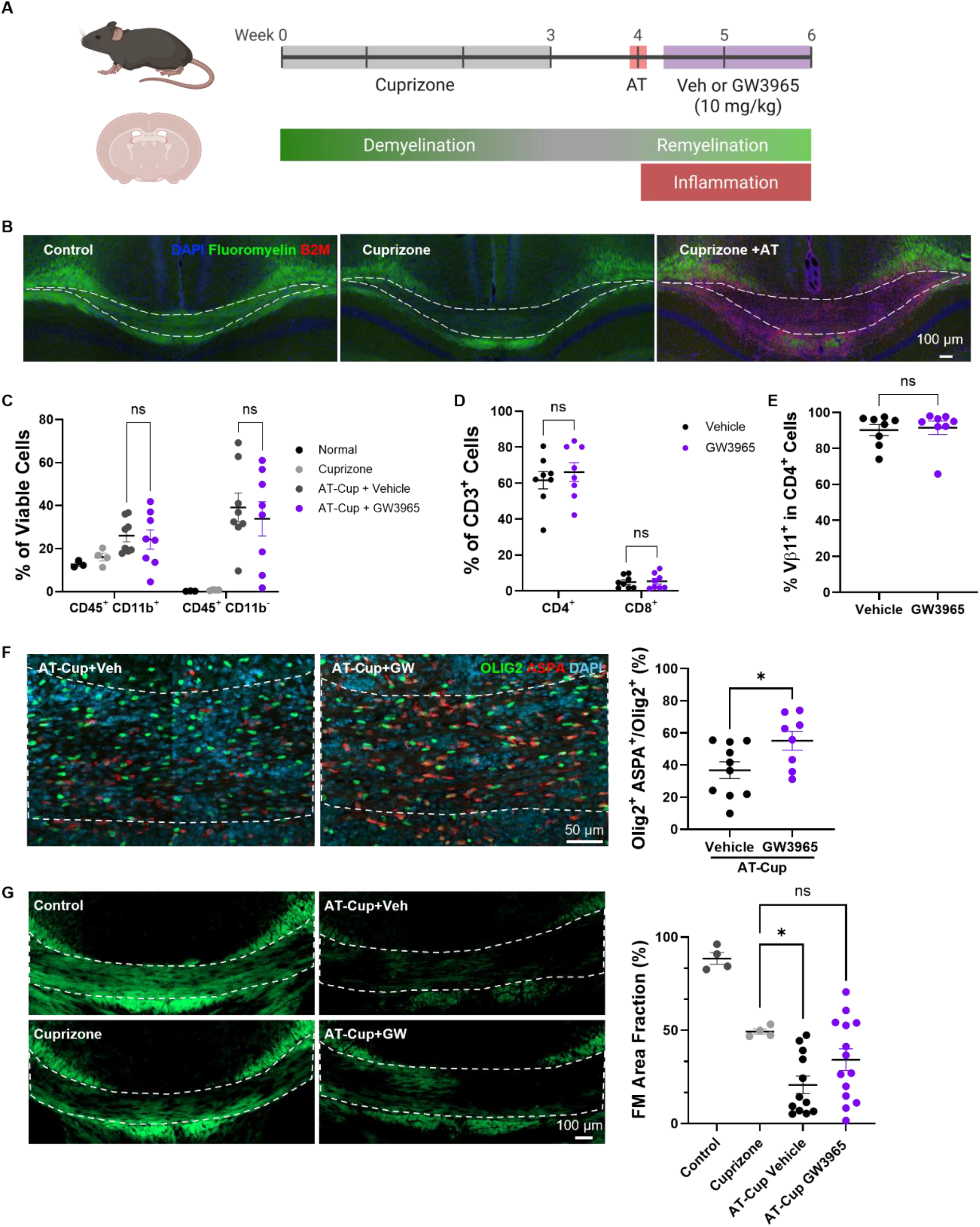
The LXR agonist GW3965 enhances OPC differentiation and augments remyelination in AT-Cup mouse model. **(A)** Schematic of experimental design. Recipient mice were fed cuprizone for 3 weeks to induce demyelination. Feed was then changed to normal chow for 1 week before adoptive transfer of Th17-polarized CD4^+^ T cells isolated from 2D2 mice. Mice were treated with vehicle or GW3965 (10 mg/kg) via daily IP injection beginning 3 days after adoptive transfer and sacrificed at 14 days post-transfer. **(B)** Representative coronal sections from the corpus callosum demonstrating accumulation of B2M-positive cells with decreased fluoromyelin staining in AT-Cup vs Cup-alone mice. **(C – E)** Flow cytometric analysis of immune cell populations within the corpus callosum. *n =* 8 mice per group for AT-Cup. **(C)** Frequency of myeloid (CD45^+^ CD11b^+^) and lymphoid (CD45^+^ CD11b^-^) cells. **(D)** Proportions of CD4^+^ and CD8^+^ T cells among CD3^+^ cells. **(E)** Proportion of Vβ11^+^ 2D2 cells among CD4^+^ T cells. **(F)** Immunofluorescence staining of corpus callosum for Olig2 and the mOL marker ASPA. Shown are representative images (*left*) and quantification of Olig2^+^ASPA^+^ mOLs (*right*). *n =* 8 mice per group. **(G)** Fluoromyelin (FM) staining of corpus callosum. Shown are representative images (*left*) and quantification of FM area fraction from control mice (*n =* 4), Cup-alone mice (*n =* 4), and AT-Cup mice treated with vehicle (*n =* 12) or GW3965 (*n =* 14). \**P* <0.05. Statistics performed using two-tailed Welch’s t-test (C, E, F), two-way ANOVA (D), or one-way ANOVA with Dunnett multiple comparison test (G). Graphs shown as mean ± SEM.

Because the LXRs are expressed in peripheral immune cells, we sought to minimize the effect of drug treatment on peripheral immune activation and CNS immune infiltration in order to isolate the impact on OPC responses to inflammation. We therefore waited until 3 days after adoptive transfer to begin treatment with GW3965 (10 mg/kg) or vehicle via daily intraperitoneal (IP) injection. As hoped, this paradigm produced minimal effects on the CNS immune infiltrate. Flow cytometry performed from corpus callosum (Supplementary Fig. 8A) found that GW3965 treatment did not change the percentage of myeloid cells or infiltrating lymphocytes (Fig. 5C and Supplementary Fig. 8B) nor the proportions of CD4^+^ and CD8^+^ T cells (Fig. 5D and Supplementary Fig. 8C). Staining for Vβ11, the T cell receptor expressed by 2D2^+^ T cells, showed the majority of infiltrating CD4^+^ T cells were the adoptively transferred Th17 cells, and no difference was observed in Vβ11^+^ CD4^+^ T cell infiltration with GW3965 treatment (Fig. 5E and Supplementary Fig. 8D). There was no change in the percentage of infiltrating Th1 or Th17 cells (Supplementary Fig. 8E), IL-17 or IFN-γ expression in CD4^+^ cells (Supplementary Fig. 8F), IFN-γ expression in CD8^+^ T cells (Supplementary Fig. 8G and H), or the percentage of effector and central memory CD4^+^ or CD8^+^ T cells (Supplementary Fig. 8I). The only difference observed following GW3965 treatment was a small but statistically significant decrease in the proportion of IL-17A^+^IFN-γ^+^ double-positive (Th17.1) CD4^+^ cells (Supplementary Fig. 8E). Lastly, GW3965 did not affect myeloid cell activation, as assessed by MHC class II expression (Supplementary Fig. 8J).

We then examined whether LXR activation with GW3965 impacted OPC differentiation and remyelination. Despite having minimal effect on peripheral immune activation and CNS immune infiltration, GW3965 treatment enhanced OPC differentiation, as assessed by expression of the mOL marker ASPA within Olig2^+^ OL-lineage cells (Fig. 5F). To measure intact myelin formation, we used fluoromyelin staining and found that treatment with GW3965 attenuated the inhibitory effect of adoptive transfer on myelin repair (Fig. 5G). These data demonstrate that pharmacologic LXR activation overcomes inflammation-induced differentiation blockade, enhancing remyelination despite the presence of an unfavorable local immune environment.

## Discussion

A growing body of evidence suggests an inhospitable environment within multiple sclerosis lesions plays a major role in remyelination failure, preventing OL maturation despite the continued presence of OPCs.^7^ Chronic neuroinflammation appears to be a primary factor, inhibiting mOL differentiation while co-opting OPCs to take on immune-like functions.^10–18^ Failure to overcome the negative influences of inflammation may underlie the disappointing clinical results of several candidate drugs that enhance OPC maturation under non-inflammatory conditions.^15,90–94^

In this study, we examined the metabolic determinants of OPC responses to inflammation. *In vitro*, exposure to IFN-γ, a potent inducer of iOLGs^11,16^ that is ubiquitously present in multiple sclerosis lesions from both lymphoid and myeloid sources,^33–37^ produced an extensive remodeling of lipid metabolism. Changes in lipid metabolism were characterized broadly by a switch toward utilization, leading to FA depletion and an increase in FAO. Similar transcriptional changes were observed in mouse models and human multiple sclerosis tissue, validating our *in vitro* system and indicating that lipid metabolic reprogramming is a conserved response to pathologic inflammatory environments *in vivo*. Critically, we found that these changes in lipid metabolism are not simply a consequence of inflammatory signals but also a driver of OPC functional responses that can be therapeutically manipulated to change OPC fate. In particular, we identified the LXRs as master regulators of lipid metabolism that can be therapeutically targeted *in vivo* to enhance myelin repair despite an unfavorable immune environment. To our knowledge, no prior treatment approaches have been shown to overcome cytokine-mediated inhibition of OPC differentiation.

The LXRs represent intriguing therapeutic targets in multiple sclerosis. Previous work found that a common allelic variant of *NR1H3* (LXRα) confers increased risk of developing progressive multiple sclerosis, with a distinct loss-of-function variant (rs61731956) associated with a severe and rapidly progressive form of multiple sclerosis in two multi-incident families.^95^ Given that progression is driven by compartmentalized responses in the CNS (including the consequences of chronic demyelination) as opposed to peripheral immune responses, the LXRs may be particularly relevant in the context of remyelination failure and other pathophysiological processes underlying progressive multiple sclerosis. Moreover, while we found beneficial effects on oligodendroglial responses, previous studies have demonstrated that pharmacologic activation of the LXRs favorably modulates microglia, enhancing their handling of myelin debris to sustain regenerative phagocytic functions.^96–98^ Similar to our findings in oligodendroglia, LXR expression in CNS macrophages and microglia increases under inflammatory conditions and in multiple sclerosis lesions,^88,99^ suggesting an adaptive response. These convergent findings suggest that LXR activation might enhance remyelination and slow progression through effects on multiple cell types within chronic multiple sclerosis lesions. Finally, several LXR agonists have already moved through clinical development.^100–104^ Although these have been limited by effects on hepatic lipogenesis and systemic hyperlipidemia, particularly through LXRα, isoform-selective and CNS-targeting strategies are in development to overcome these challenges.^100^

A critical role for FA and lipid metabolism in modulating the inflammatory versus repair functions of OPCs should perhaps not be surprising. OPC maturation requires a lipogenic state and sufficient FAs to produce the abundant and diverse phospholipids that constitute myelin membranes.^105^ Conversely, lipid catabolic pathways such as lipolysis and FAO have been implicated in immune cell activation. For instance, CD8^+^ T cells rely on lipolysis and FAO to become memory cells,^106,107^ and FAO is critical for natural killer (NK) cell function.^108^ Thus, the transition from a lipogenic to a lipid catabolic state is well-positioned as a metabolic switch in OPCs favoring immune-like functions over maturation to myelinating OLs.

Several of our findings merit further study. For instance, we found that LXR activation promoted triglyceride synthesis, which contributed to its functional effects in OPCs. Neutral lipid accumulation has been shown to play both adaptive and maladaptive roles in various cell types,^80^ so further investigation will be necessary to understand the purpose, precise localization, and subsequent fate of triglycerides in iOLGs. Additionally, the unique role of oleic acid (monounsaturated) versus palmitic acid (saturated) supplementation merits further study to understand their divergent effects on iOLGs. In this respect, it is intriguing that expression of *Scd1* and *Scd2*, the enzymes responsible for conversion of saturated to monounsaturated fatty acids, was decreased in oligodendroglia under inflammatory conditions and increased by LXR activation.

In conclusion, we identified lipid metabolism as a critical regulator of OPC responses to inflammation, controlling the fate decision between iOLG formation and maturation into a myelinating OL. Importantly, we found that lipid metabolic pathways represent druggable targets for overcoming the inhibitory effect of inflammation on myelin repair, a currently intractable obstacle to remyelinating therapies in multiple sclerosis. Our findings suggest that targeted LXR agonists hold particular promise as candidate remyelinating agents.

## Data availability

All data supporting the findings of this study are presented and/or tabulated within the article or supplementary materials. RNAseq data has been deposited in the Gene Expression Omnibus (GEO) database and will be made publicly available upon publication. No unique code was generated.

## Acknowledgements

The authors thank Yasmin Resto and Danny Galleguillos for their support and advice. We thank Dr. Ira Schulman for kindly sharing LXR knockout mice. Fig. 1A, 2D, 5A, and Supplementary Fig. 8A were created using BioRender.

## Supplementary material

Supplementary material includes: Supplementary Figures 1 through 8 and supplemental file S1.

