## Supplementary Figures for "Remodeling oligodendrocyte lipid metabolism via liver X receptors overcomes inflammatory blockade of remyelination"

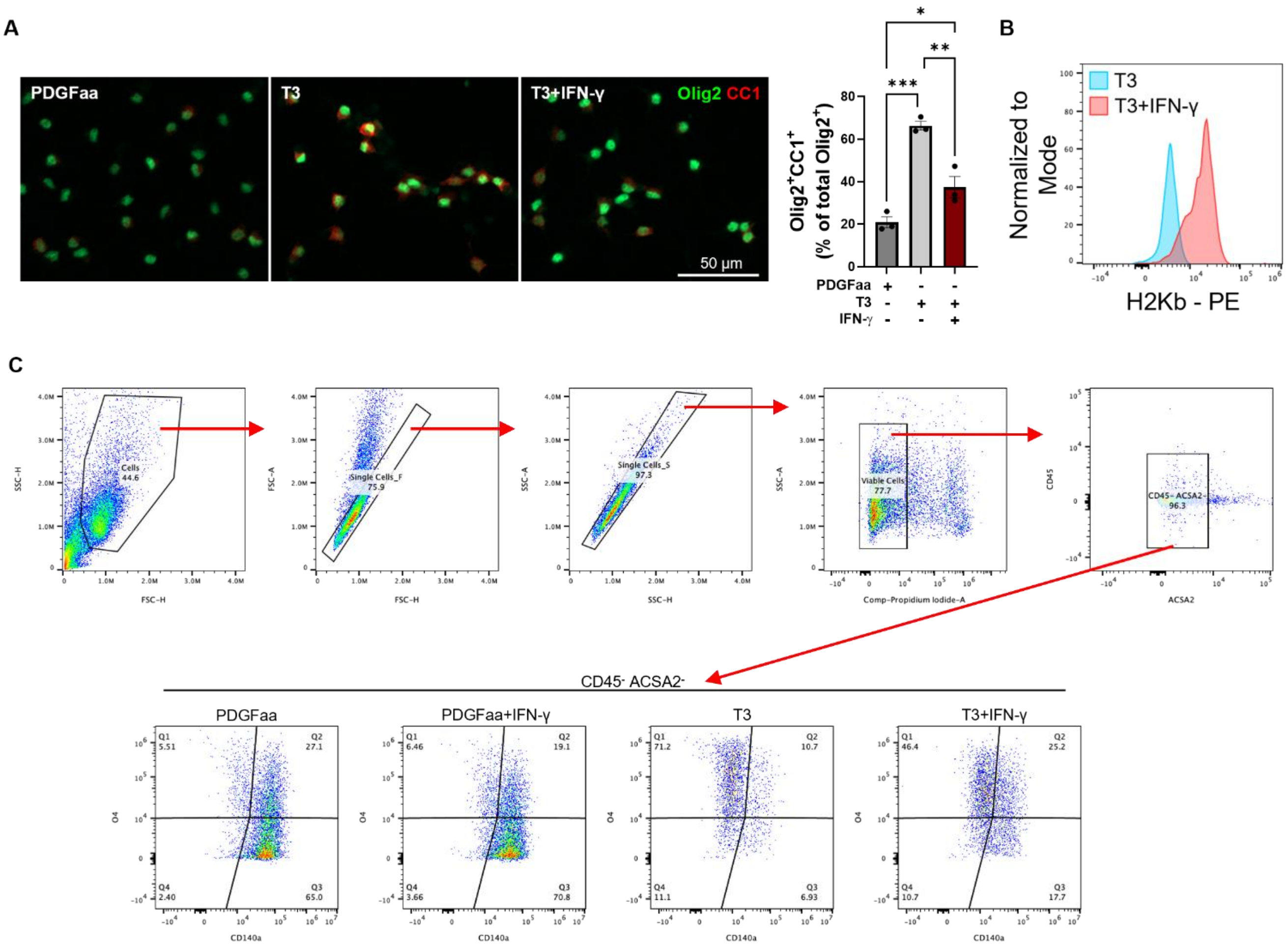


**Supplementary Figure 1. Induction of mouse iOLGs with IFN-γ**. Corresponding to Figure 1A-C. Primary mouse OPCs were cultured in PDGFaa (to maintain a proliferative state) or T3 (to induce maturation) ± IFN-γ for 48 h. **(A)** OPC maturation was assessed by immunofluorescent staining for Olig2 and CC1. (*Left*) Representative images and (*right*) quantification from $n = 3$ independent experiments per group. Each experiment consisted of independent biological samples with quantification from four fields per sample. Mean ± SEM. **(B)** Representative histogram of H2Kb (MHC class I) expression in iOLGs, measured via flow cytometry in CD140a$^-$ O4$^+$ population. **(C)** Gating strategy for flow cytometric analysis of iOLGs. *$P$ <0.05, **$P$<0.01, ***$P$<0.001 by one-way ANOVA with Tukey multiple comparisons test.

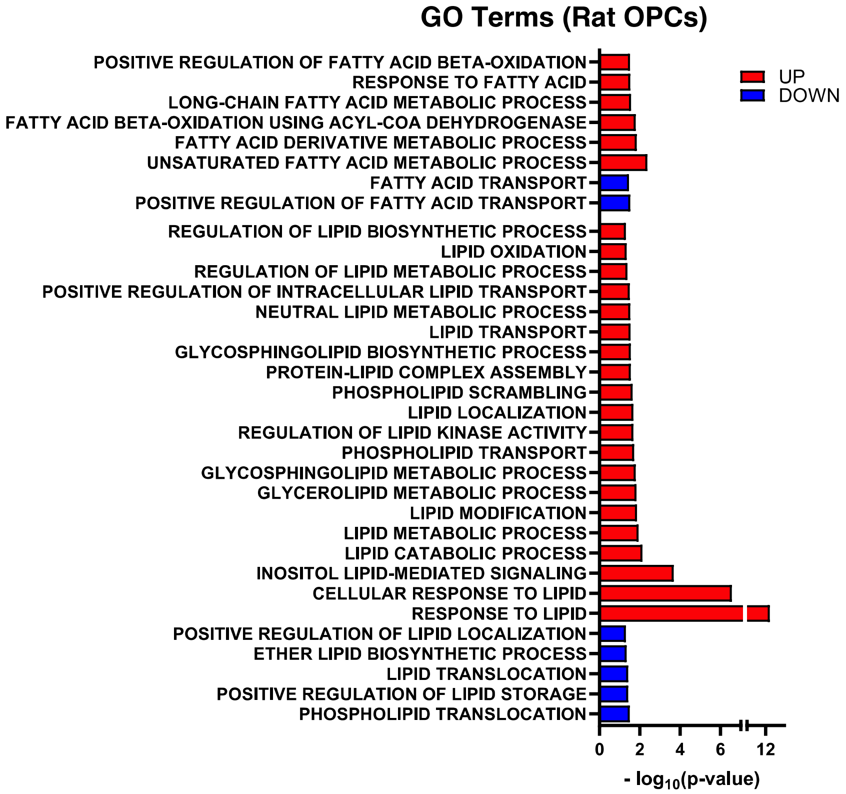


**Supplementary Figure 2. Rat iOLGs reprogram lipid metabolism.** RNAseq was performed from rat OPCs treated with T3 or T3+IFN-γ for 24 h. Shown are GO terms associated with fatty acid and lipid metabolism that were significantly upregulated (red) or downregulated (blue) with IFN-γ treatment. *n* = 3 biological replicates. P-values were calculated using a hypergeometric over-representation test on the set of differentially expressed genes.

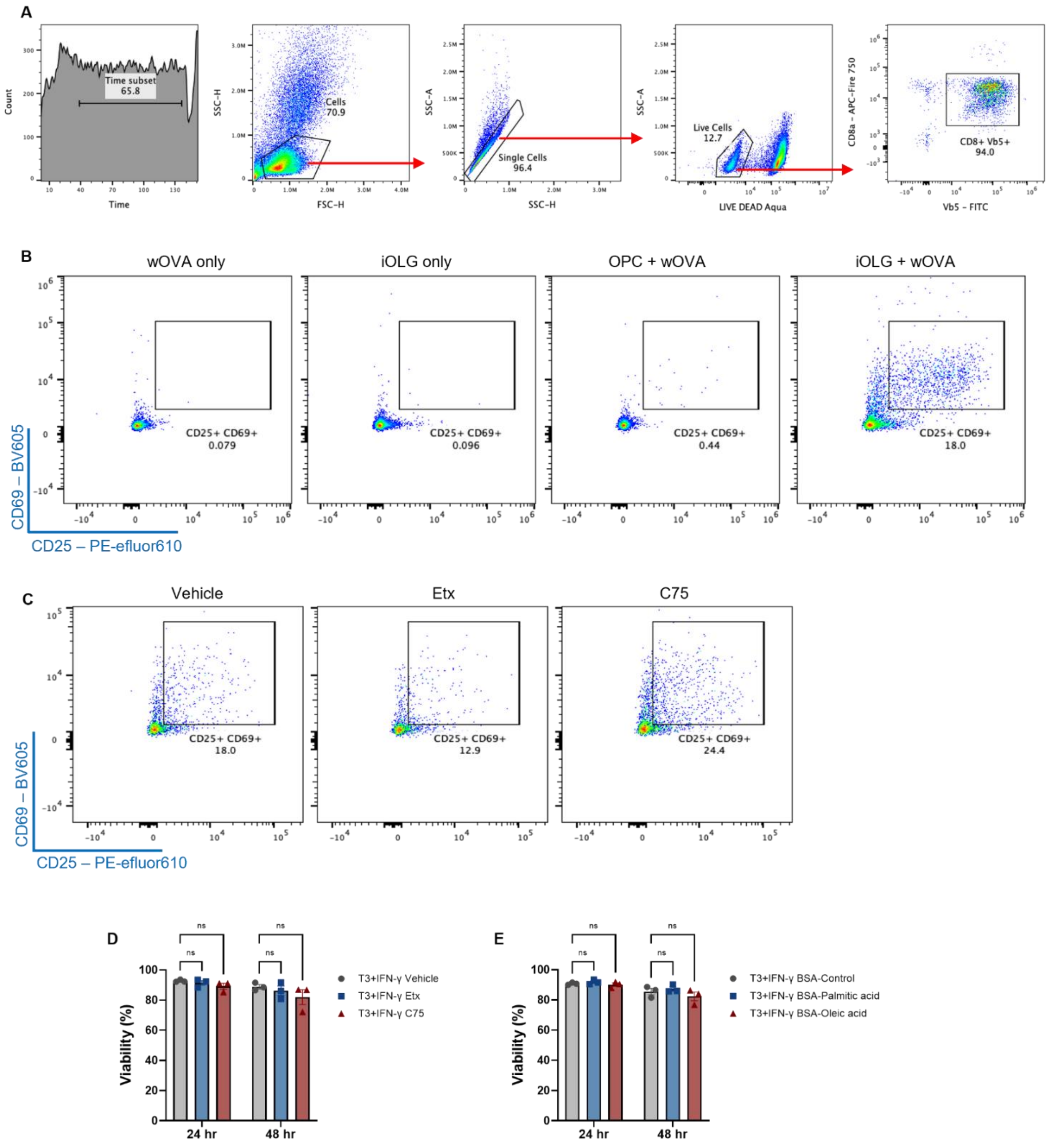


**Supplementary Figure 3. iOLG-CD8$^+$ T cell co-culture assays. (A)** Gating strategy for flow cytometric analysis of CD8$^+$ Vβ5$^+$ T cells following co-culture with iOLGs. **(B)** Representative flow plots of CD25 and CD69 expression in CD8$^+$ T cells cultured under different treatment conditions. CD8$^+$ T cell activation occurred only when OPCs were treated with both IFN-γ and wOVA, indicating that wOVA must be processed into peptides and cross-presented by iOLGs for CD8$^+$ T cell activation. **(C)** Representative flow plots of CD8$^+$ T cells activated by IFN-γ-induced iOLGs treated with vehicle, Etx (40 μM), or C75 (20 μM). **(D and E)** Live cell imaging with Incucyte was performed to measure viability of iOLGs cultured in T3+IFN-γ ± Etx, C75, or FAs for 48 h. $n = 3$ independent experiments performed in triplicate. Statistical analysis was performed using one-way ANOVA with Dunnett multiple comparisons test. ns = not significant.

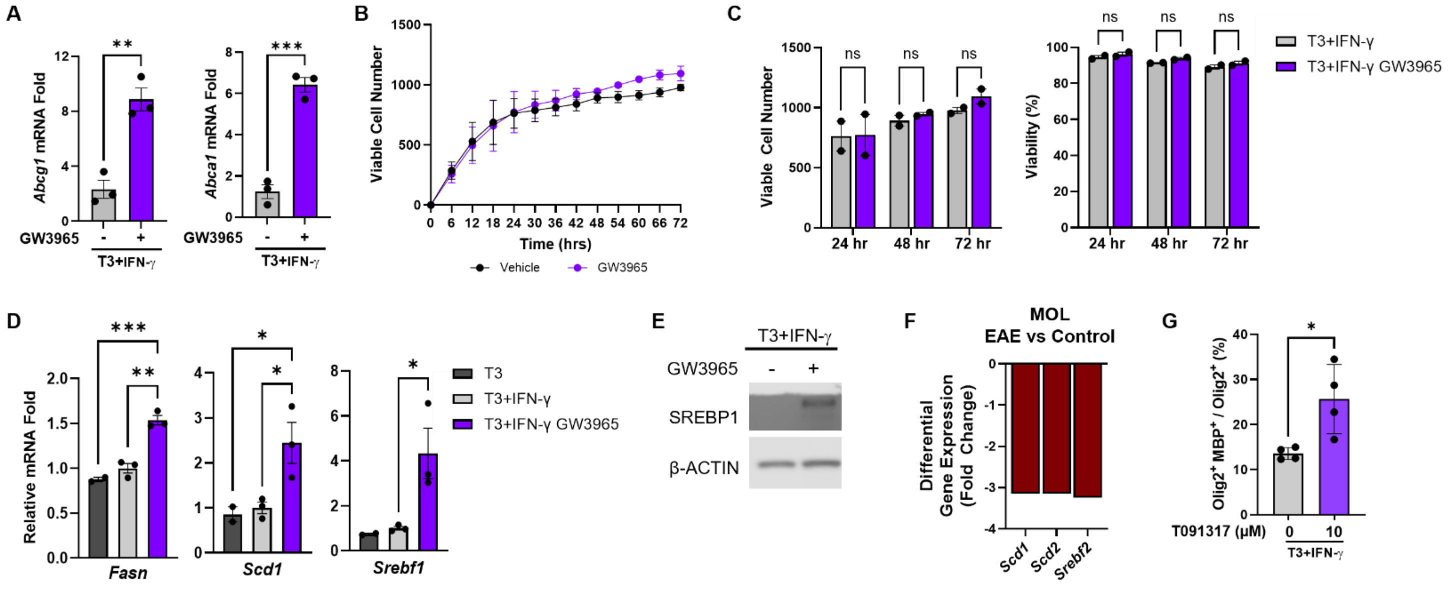


**Supplementary Figure 4. GW3965 activates LXR in mouse OPCs without impacting cell proliferation or viability**. **(A)** Expression of LXR target genes *Abcg1* and *Abca1* in mouse OPCs treated with T3+IFN-γ ± 20 µM GW3965, measured by qRT-PCR. *n* = 3 independent experiments performed in triplicate. **(B and C)** Live cell imaging with Incucyte was performed to measure cell proliferation and viability in OPCs cultured in T3+IFN-γ ± 20 µM GW3965 for 72 h. *n* = 2 independent experiments performed in triplicate. **(D)** mRNA expression was measured via qRT-PCR from mouse OPCs treated with T3, T3+IFN-γ, or T3+IFN-γ+GW3965 (20 µM) for 24 hours. *n* = 3 independent experiments performed in triplicate. **(E)** Representative Western blot demonstrating induction of SREBP1 (*Srebf1*) protein expression in mouse OPCs treated for 48 h with GW3965 (20 µM). **(F)** Differential expression of *Scd1*, *Scd2*, and *Srebf2* in MOLs from EAE versus control mice. From Falcão, et al. **(G)** Mouse OPCs were cultured in T3+IFN-γ ± the structurally distinct LXR agonist T091317 (10 µM) for 72 h followed by immunofluorescent staining for Olig2 and MBP. *n* = 4 independent experiments. Each experiment consisted of independent biological samples with quantification from four fields per sample. *$P$ <0.05, **$P$<0.01, ***$P$<0.001 by two-tailed Welch's *t* test (A, G), two-way ANOVA (C), or one-way ANOVA with Tukey multiple comparisons test (D). Graphs shown as mean ± SEM.

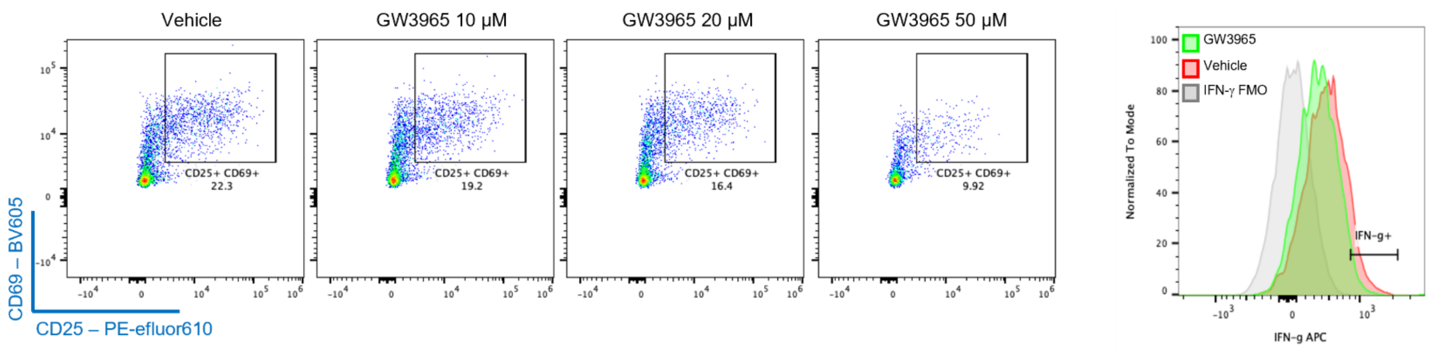


**Supplementary Figure 5. LXR activation suppresses iOLG-mediated CD8$^{+}$ T cell activation**. Representative flow plots of CD25 and CD69 expression (*left*) and representative histogram of IFN-γ expression (*right*) in CD8$^{+}$ T cells co-cultured with wOVA-loaded mouse iOLGs that had been treated with the indicated doses of GW3965.

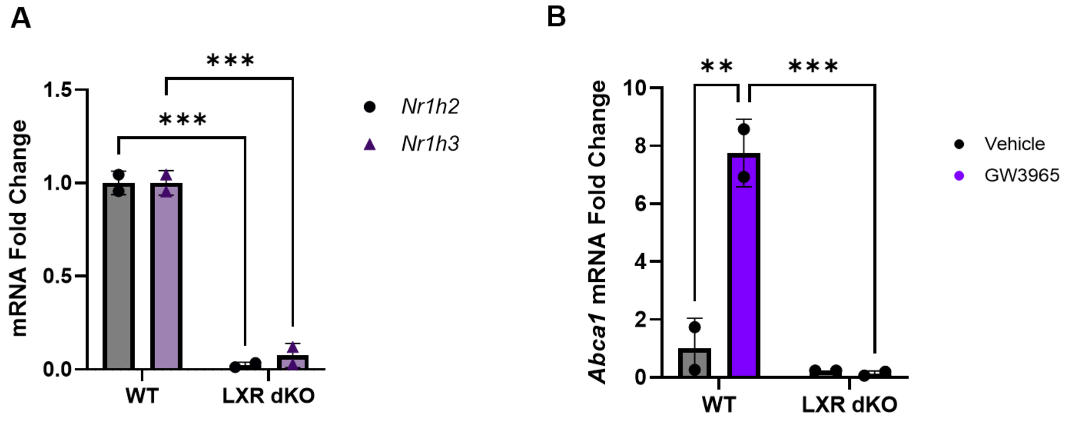


**Supplementary Figure 6. Validation of LXR dKO OPCs**. **(A)** Expression of both LXR isoforms (*Nr1h2* and *Nr1h3*) was eliminated in LXR dKO OPCs, as assessed by qRT-PCR. $n = 2$ independent experiments performed in triplicate; mean ± SEM. **(B)** The LXR agonist GW3965 (20 μM) failed to induce expression of the LXR target gene *Abca1* in LXR dKO OPCs. $n = 2$ independent experiments performed in triplicate; mean ± SEM. ***P*<0.01, ****P*<0.001 by two-way ANOVA.

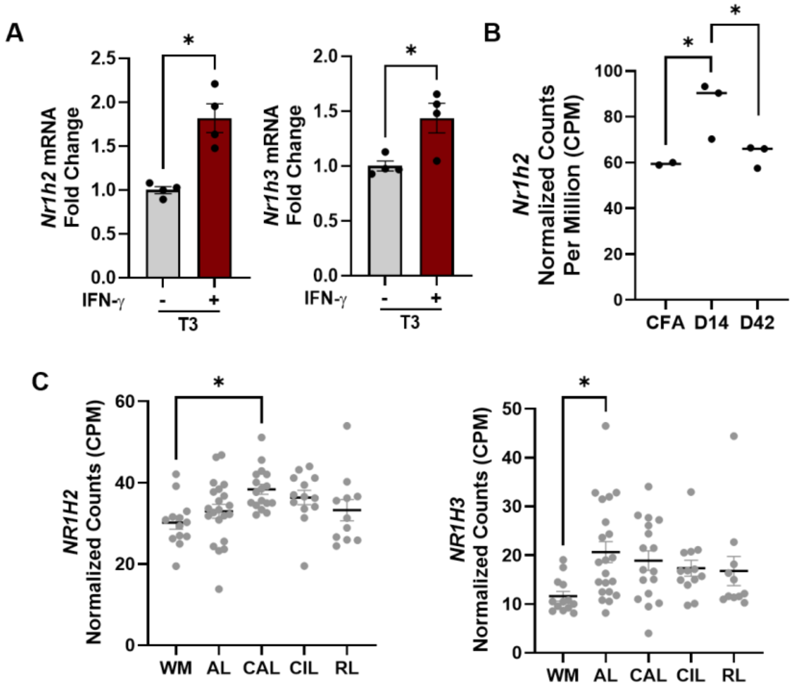


**Supplementary Figure 7. Increased LXR expression in oligodendroglia under inflammatory conditions *in vitro* and *in vivo*. (A)** mRNA expression was measured via qRT-PCR in mouse OPCs treated with T3 or T3+IFN-γ for 24 h. $n = 4$ independent experiments performed in triplicate; mean ± SEM. **(B)** Expression of LXRβ (*Nr1h2*) in pseudobulked OL-lineage cells from CFA-alone, EAE Day 14, and EAE Day 42 mice using snRNAseq data from Mace and Gadani, et al. **(C)** LXRβ (*NR1H2*) and LXRα (*NR1H3*) expression was examined in OLs from different MS lesion subtypes: white matter (WM) from healthy controls, active demyelinated lesions (AL), chronic active demyelinated lesions (CAL), chronic inactive demyelinated lesions (CIL), and remyelinated lesions (RL) from MS tissue. Analysis of snRNAseq data from Macnair, et al. **P* <0.05 by two-tailed Welch's t-test (A), Wald test adjusted for FDR (B), or glmmTMB *FDR < 0.05 (C).

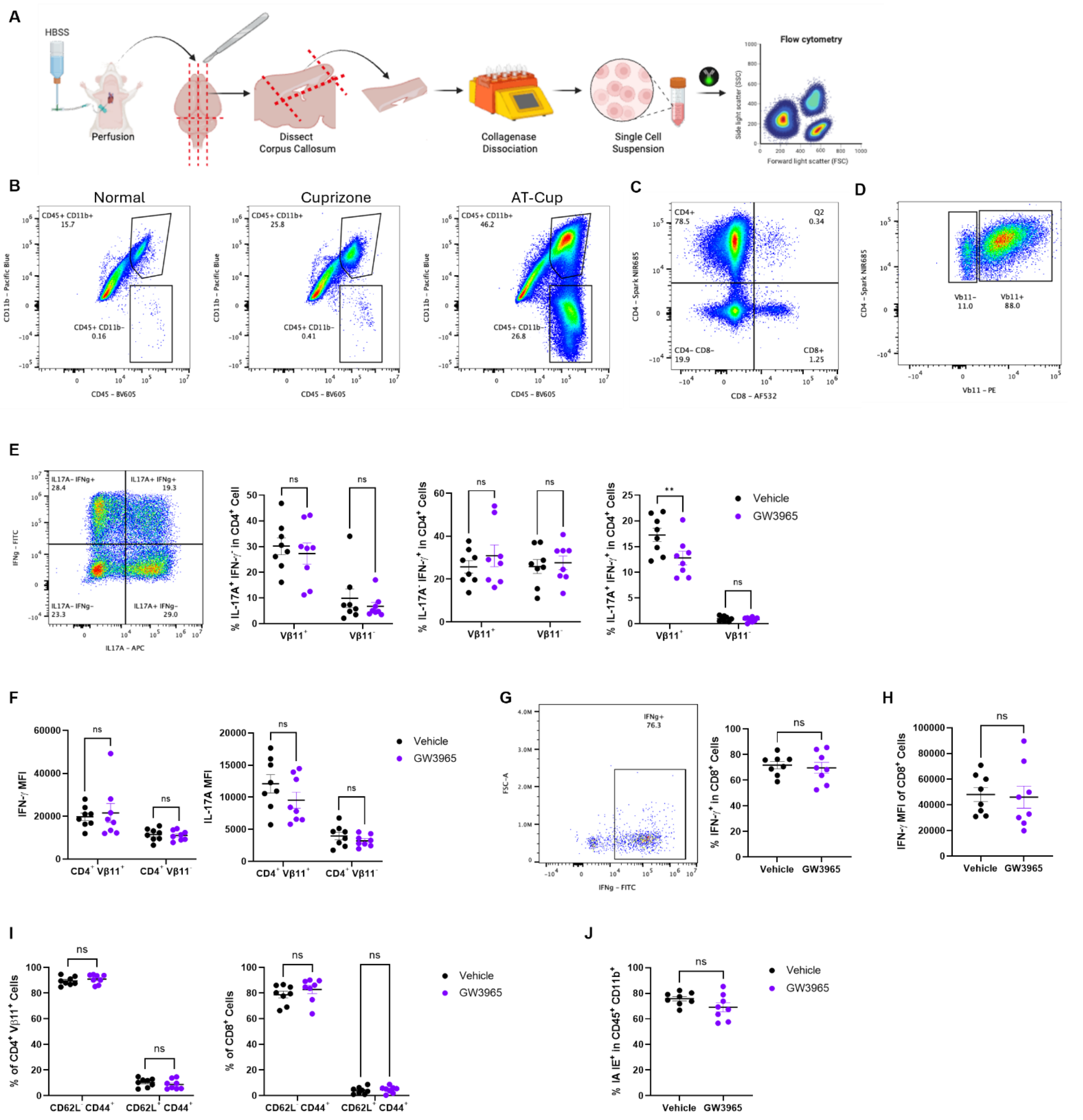


**Supplementary Figure 8. Minimal effects of GW3965 treatment on the CNS immune infiltrate in the AT-Cup mouse model. (A)** Schematic of workflow for flow cytometric analysis of corpus callosum. Mice were perfused with cold HBSS$^{+/+}$ and the corpus callosum was dissected. Tissue was dissociated with collagenase, and debris was removed prior to antibody staining. **(B)** Representative flow plots of CD45$^{+}$ CD11b$^{+}$ (myeloid) and CD45$^{+}$ CD11b$^{-}$ (lymphoid) cells in the corpus callosum from control, cuprizone-alone, and AT-Cup mice. **(C)** Representative flow plot of CD4$^{+}$ and CD8$^{+}$ T cells gated on CD3$^{+}$ cells. **(D)** Representative flow plot of Vβ11 gating. **(E)** (*Left*) Representative flow plot of IL-17A and IFN-γ expression in CD4$^{+}$ T cells. (*Right*) Quantification of Th17 (IL-17A$^+$ IFN-γ$^-$), Th1 (IL-17A$^-$ IFN-γ$^+$), and Th17.1 (IL-17A$^+$ IFN-γ$^+$) CD4$^+$ T cells. **(F)** Mean fluorescence intensity (MFI) of IFN-γ and IL-17A expression in CD4$^+$ Vβ11$^+$ or CD4$^+$ Vβ11$^-$ cells. **(G)** Representative flow cytometry plot (*left*) and quantification of IFN-γ$^+$ frequency (*right*) among CD8$^+$ T cells. **(H)** MFI of IFN-γ expression in CD8$^+$ T cells. **(I)** Frequencies of CD62L$^-$ CD44$^+$ and CD62L$^+$ CD44$^+$ cells among CD4$^+$ Vβ11$^+$ or CD8$^+$ T cells. **(J)** Frequencies of IA IE$^+$ (MHC class II) cells among CD45$^+$ CD11b$^+$ myeloid cells. Data presented as mean ± SEM. Each dot represents one mouse ($n$ = 8 per group). Significance was determined by two-tailed Welch's $t$ test or two-way ANOVA. **$P$<0.01.
